# Cleft lip and cleft palate (CL/P) in *Esrp1* KO mice is associated with alterations in Wnt signaling and epithelial-mesenchymal crosstalk

**DOI:** 10.1101/2019.12.12.874636

**Authors:** SungKyoung Lee, Matthew J. Sears, Zijun Zhang, Hong Li, Imad Salhab, Philippe Krebs, Yi Xing, Hyun-Duck Nah, Trevor Williams, Russ P. Carstens

## Abstract

Cleft lip is one of the most highly prevalent birth defects in human patients. However, there remain a limited number of mouse models of cleft lip and thus much work is needed to further characterize genes and mechanisms that lead to this disorder. It is well established that crosstalk between epithelial and mesenchymal cells underlies formation of the face and palate, yet the basic molecular events mediating this crosstalk are still poorly understood. We previously demonstrated that mice with ablation of the epithelial-specific splicing factor Esrp1 have fully penetrant bilateral CL/P. In this study we further investigated the mechanisms by which ablation of Esrp1 leads to cleft lip as well as cleft palate. These studies included a detailed analysis of the changes in splicing and total gene expression in embryonic ectoderm during formation of the face as well as gene expression changes in adjacent mesenchyme. We identified altered expression in components of pathways previously implicated in cleft lip and/or palate, including numerous components of the Wnt signaling pathway. These findings illustrate that maintenance of an Esrp1 regulated epithelial splicing program is essential for face development through regulation of key signaling pathways.

## INTRODUCTION

Cleft lip and/or palate (CL/P) are among the most common congenital birth defects, affecting approximately 1 in 700 live births and affected children face a variety of health and psychosocial problems as well as a need for extensive surgical and dental treatments (Dixon et al., 2011). The causes of CL/P are heterogeneous and include a variety of environmental and genetic factors. Although CL/P can be a component of disease syndromes, most cases of CL/P are non-syndromic. Cleft lip with or without cleft palate (CL/P) is more common in human patients than isolated cleft palate (CP, or CPO) and these disorders are largely genetically and etiologically distinct (Fraser, 1970; Gritli-Linde, 2008).The proper development of the lip and palate is similar between humans and mice and thus mouse models have served an important role in studies to identify genes and characterize pathways that, when disrupted, lead to CL/P or CPO (Gritli-Linde, 2012; Juriloff and Harris, 2008). In mice, formation of the face commences around E9.5 and involves the five facial prominences consisting of mostly neural crest-derived mesenchyme and overlying epithelium; the frontonasal prominence (FNP) and the paired maxillary and mandibular prominences (MXP and MdP) (Jiang et al., 2006). The FNP gives rise to the lateral and medial nasal prominences (LNP and MNP) and these prominences grow into close apposition and by E12.5 the nasal and maxillary prominences fuse to form the upper lip and primary palate. Defects in the growth and/or fusion of these prominences result in CL/P. The formation of the secondary palate is a separate developmental process that occurs from E12-E15.5 when the palatal shelves emerge from the maxillary prominences, elevate, and fuse in the midline (Jiang et al., 2006). Defects in any of these steps can lead to isolated cleft palate (CP). While CL/P is the more common human clinical presentation, there are many mouse models of CPO, yet relatively few for CL/P (Gritli-Linde, 2008). Thus, while studies of the cellular and molecular changes that occur in developing mouse face between E9.5-12.5 have the greatest relevance to the pathogenesis of CL/P, newer mouse models for CL/P are needed to further define genes and pathways involved in CL/P pathogenesis.

During craniofacial development, the ectoderm and derivative epithelial cells of the facial prominences and palate provide signals required for mesenchymal proliferation and patterning. At the same time the mesenchyme provides feedback to epithelial cells and these reciprocal epithelial-mesenchymal interactions are crucial for normal facial and palatal development (Jiang et al., 2006; Wedden, 1987). These interactions involve signaling pathways for the Wnt, Bmp/TGF-β, Hedgehog, and Fgf families and mutations in components of these signaling pathways have been shown to cause CL/P in human patients (Reynolds et al., 2019).

Our lab identified the Epithelial Splicing Regulatory Proteins 1 and 2 (ESRP1 and ESRP2) as exquisitely epithelial-specific regulators of multiple target transcripts, including an event in fibroblast growth factor receptor 2 (Fgfr2) whose dysregulated splicing is associated with cleft palate (Bebee et al., 2015; Rice et al., 2004; Warzecha et al., 2009a). While there is some functional redundancy between these two paralogous proteins, only ESRP1 is essential, since neither depletion nor ablation of ESRP2 alone leads to significant splicing alterations and Esrp2 KO mice have no apparent phenotype (Bebee et al., 2015; Warzecha et al., 2009b). In contrast, the loss of ESRP1 alone can lead to substantial alterations in splicing of numerous target transcripts, albeit the loss of both ESRP1 and ESRP2 is generally associated with larger changes in splicing in larger sets of transcripts. We previously showed that ablation of Esrp1 alone in mice led to fully penetrant bilateral CL/P, adding this splicing factor to the limited number of genes whose ablation leads to this defect in mice. We therefore hypothesized that facial development is dependent on Esrp1-regulated splicing events and that further studies using these mice have the potential to reveal novel molecular mechanisms and signaling pathways whose dysregulation leads to CL/P. We carried out a more detailed analysis of the defects in facial and palatal development and an extensive analysis of changes in alternative splicing in the epithelial cells of the facial prominences as well as changes in total gene expression - in both epithelial cells and underlying mesenchyme. We identified reduced expression of several genes in *Esrp1* KO ectoderm, including several canonical Wnts, including Wnt9b whose deletion has previously been shown to lead to CL/P in mice (Carroll et al., 2005; Ferretti et al., 2011; Jin et al., 2012).These changes in expression of Wnts in ectoderm were accompanied by reduced expression of canonical Wnt targets genes and reduced proliferation in adjacent mesenchyme. These observations indicate that Esrp1 plays an important role in epithelial-mesenchymal cross-talk during craniofacial development. We also noted a defect in epithelial cell fusion during lip formation and in palatal explant cultures indicating that Esrp1 is required for two distinct processes, growth and fusion, that are required for proper lip and palatal formation.

## RESULTS

### Condition ablation of *Esrp1* in surface ectoderm leads to CL/P

We examined the tissue specificity of ESRP1 action via generation of new mouse genetic models. First, we generated mice with endogenously FLAG epitope tagged ESRP1 that enabled ESRP1 protein expression to be tracked with high precision and sensitivity. Analysis of these mice (Esrp1^FLAG/FLAG^ mice, Fig. S1) confirmed that ESRP1 protein is specifically expressed in surface ectoderm at E11.5 as well as epithelial cells of the secondary palate as was previously described for *Esrp1* mRNA (Revil and Jerome-Majewska, 2013; Warzecha et al., 2009a). Second, we generated mice with conditional ablation of *Esrp1* in surface ectoderm derived from *Esrp1*^*flox/flox*^ mice and transgenic *Crect* mice that express Cre specifically in surface ectoderm and derivatives starting at E8.5, a time point prior to lip fusion (Reid et al., 2011). Compared to littermate controls, we noted bilateral CL/P in *Esrp1*^*flox/flox*^; *Crect*^+/−^embryos at E18.5 (Fig. 1A). While the phenotype was less extensive than we previously observed in *Esrp1*^−/−^ mice, there was a clear cleft of the primary palate and failure of the lip processes to come together to form a midline philtrum. To further characterize the CL/P defect we also performed staining for bone and cartilage structures which confirmed a completely cleft secondary palate and a bilateral cleft of the primary palate. The palatine bones (p) were hypoplastic and located to the side and the palatal processes of the maxilla (ppmx) were similarly dysmorphic. The premaxilla (pm) was undeveloped with a poor connection to the maxilla (m) and thus extended out in front of the face. (Fig. 1B). These findings confirm that ESRP1 functions specifically in the ectoderm and its derivatives, but that the resultant changes affect the patterning of the underlying mesenchyme.

**Fig. 1.**
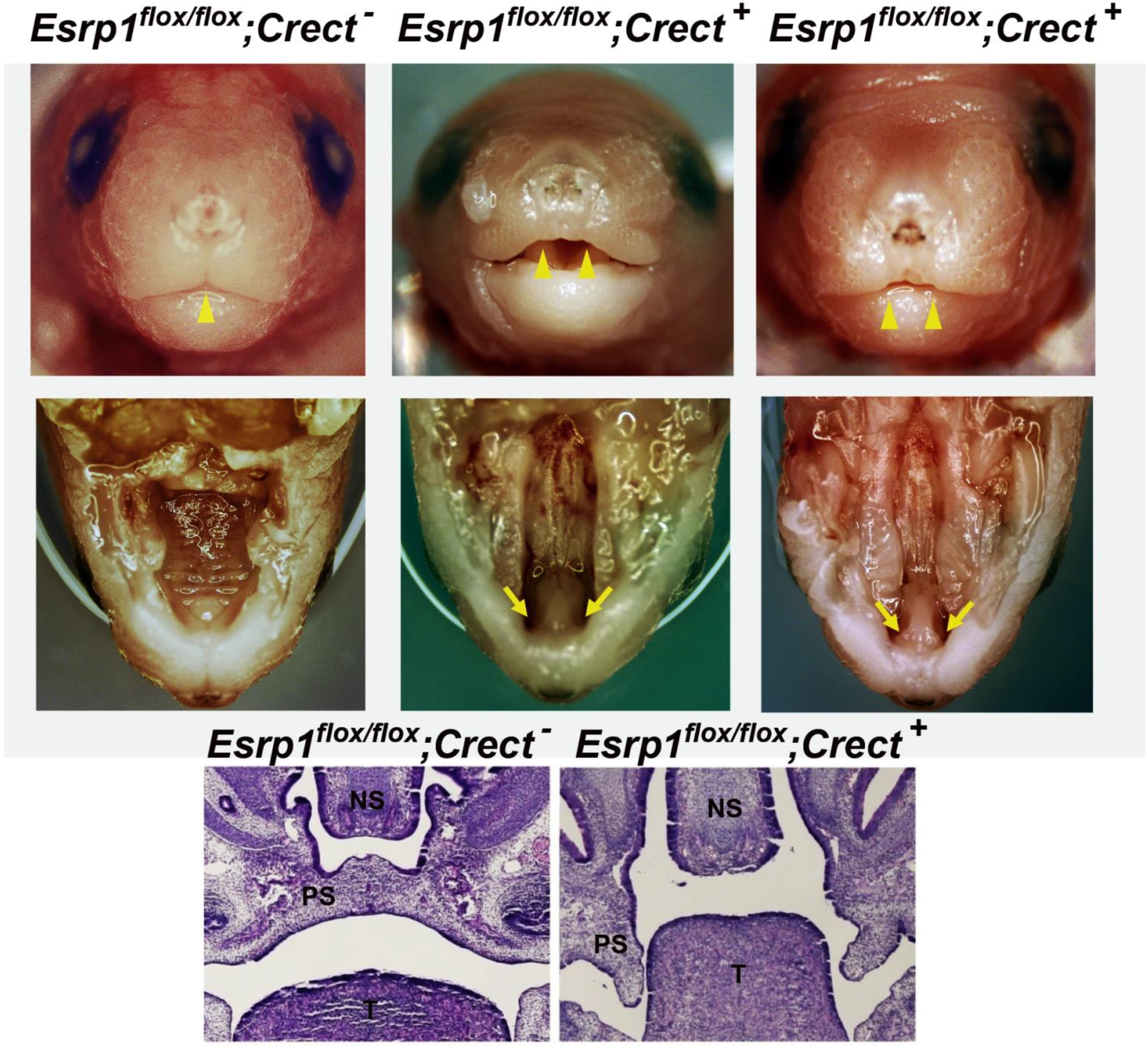
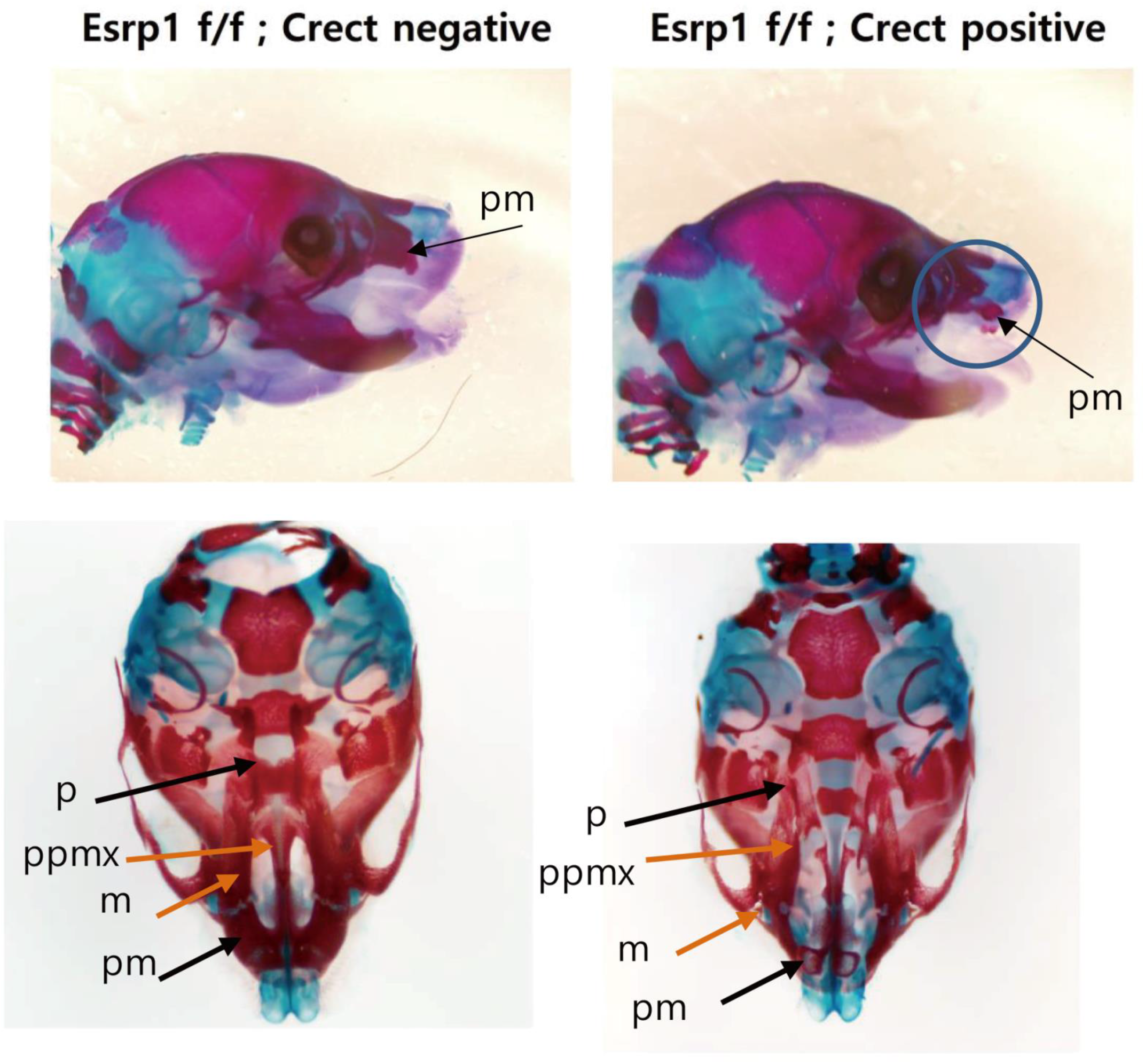
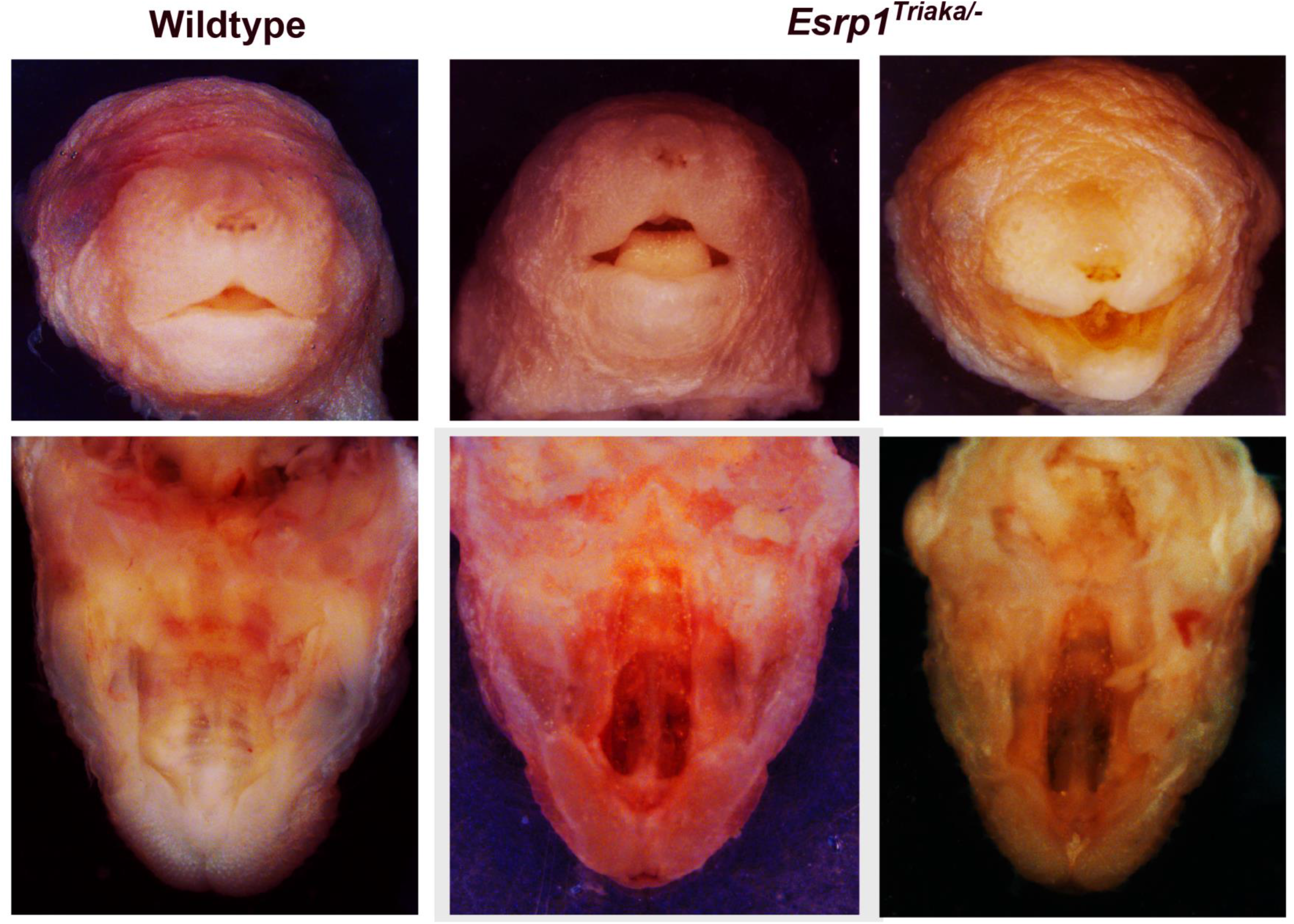
Conditional ablation of *Esrp1* in surface ectoderm leads to CL/P. (A) Frontal views of control (*Esrp1*^*flox/flox*^) mice and two *Esrp1*^*flox/flox*^; *Crect*^+/−^ cKO E18.5 embryos showing that the midline philtrum observed in control mice (single arrowhead) is absent in cKO mice as the upper lip primordia fail to meet at the midline (double arrowheads). Views of the palate after removal of the mandible show clefting of the secondary palate as well as the primary palate in the cKO mutants (arrows). H & E images of coronal sections show that compared to the normal palate formed in control embryos, the palatal processes in *Esrp1*^*flox/flox*^; *Crect*^+/−^ embryos are hypoplastic and do not elevate. PS, palatal shelf; NS, Nasal Septum; T, Tongue. (B) Alizarin Red/Alcian Blue bone and cartilage stains of E18.5 embryos from the side (top) and from the ventral side with mandible removed (bottom). Defects are apparent in the cKO maxilla (m), palatine (p), premaxilla (pm), and palatal process of the maxilla (ppmx). (C) Frontal and ventral view with lower jaw removed of Esrp1^Triaka/−^ mice showing normal lip formation with midline philtrum, but cleft secondary palate.

Previous studies have shown that mice homozygous for a null mutation in Esrp1 have of the lip, primary palate and secondary palate. However, it was recently demonstrated that mice with an ENU-induced point mutation in a highly conserved region of Esrp1 (*Triaka* mice) did not show craniofacial defects, but instead show altered intestinal function, raising the issue of how sensitive facial development is to ESRP1 activity (Mager et al., 2017). In crosses between *Esrp1*^+/−^ and *Triaka* mice to generate compound mutant mice it was revealed that whereas neither *Esrp1*^+/−^ mice, nor *Triaka* homozygous animals showed overt craniofacial defects, we observed that *Esrp1*^*triaka/−*^ pups had clefting of the secondary soft and hard palate, but no apparent cleft of the lip or primary palate (Fig 1C). These data show that this null/hypomorph allelic combination is able to generate a model of cleft secondary palate in the absence of cleft primary palate, indicating that these two developmental processes are not linked, but have different sensitivities to the levels of active ESRP1. In any event, the observation of a cleft secondary palate in the absence of cleft lip in *Esrp1*^*triaka/−*^ pups strongly indicated that *Esrp1* deficiency independently leads to a failure in palate formation.

### Esrp1 ablation leads to reduced outgrowth of facial prominences associated with reduced epithelial and mesenchymal cell proliferation

Lip formation results from outgrowth and fusion of the MxP, LNP, and MNP and failure in either process can lead to cleft lip. We assessed whether reduced cell proliferation or increased apoptosis within these processes prior to lip fusion might at least partially underlie orofacial clefting in *Esrp1*^−/−^ mice. Ki67 staining of MNP and LNP in sections from E10.5 embryos showed that there was a reduction in cell proliferation in both epithelial and mesenchymal cells associated with less apparent growth of the two processes towards apposition (Fig. 2A). Evaluation of apoptosis using staining for activated caspase 3 showed no apparent differences between control and knockout embryos other than the expected apoptosis in sections where these MNP and LNP processes were beginning to fuse in control embryos. In *Esrp1*^−/−^ mice these processes either did not make contact, or if they did make contact there was no observed fusion and no apparent apoptosis at the sites of contact (Fig. 2B). In one case where we observed contact between the MNP and LNP in *Esrp1*^−/−^ mice, we further noted that there was persistent expression of E-Cadherin that was not observed in wild-type (WT) embryos (Fig 2C). We conclude that a reduction in proliferation of facial prominences contributes to CL/P in *Esrp1*^−/−^ mice, but that there is also a defect in fusion when these processes do make contact. Nonetheless, the reduction in cell proliferation in mesenchyme adjacent to *Esrp1* ablated epithelial cells indicated that there was a disruption in a communication pathway from epithelium to mesenchyme during facial development.

**Fig. 2.**
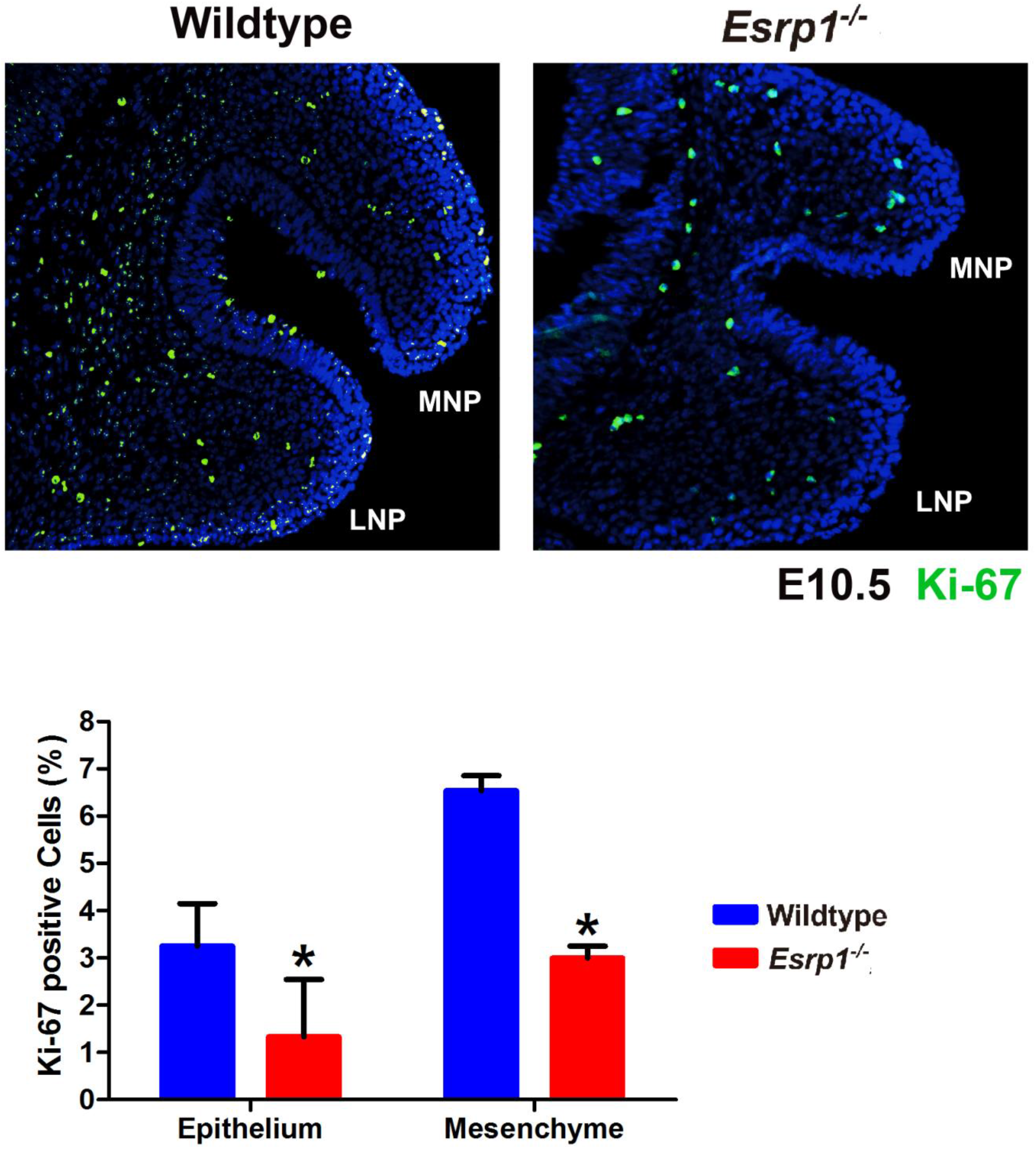
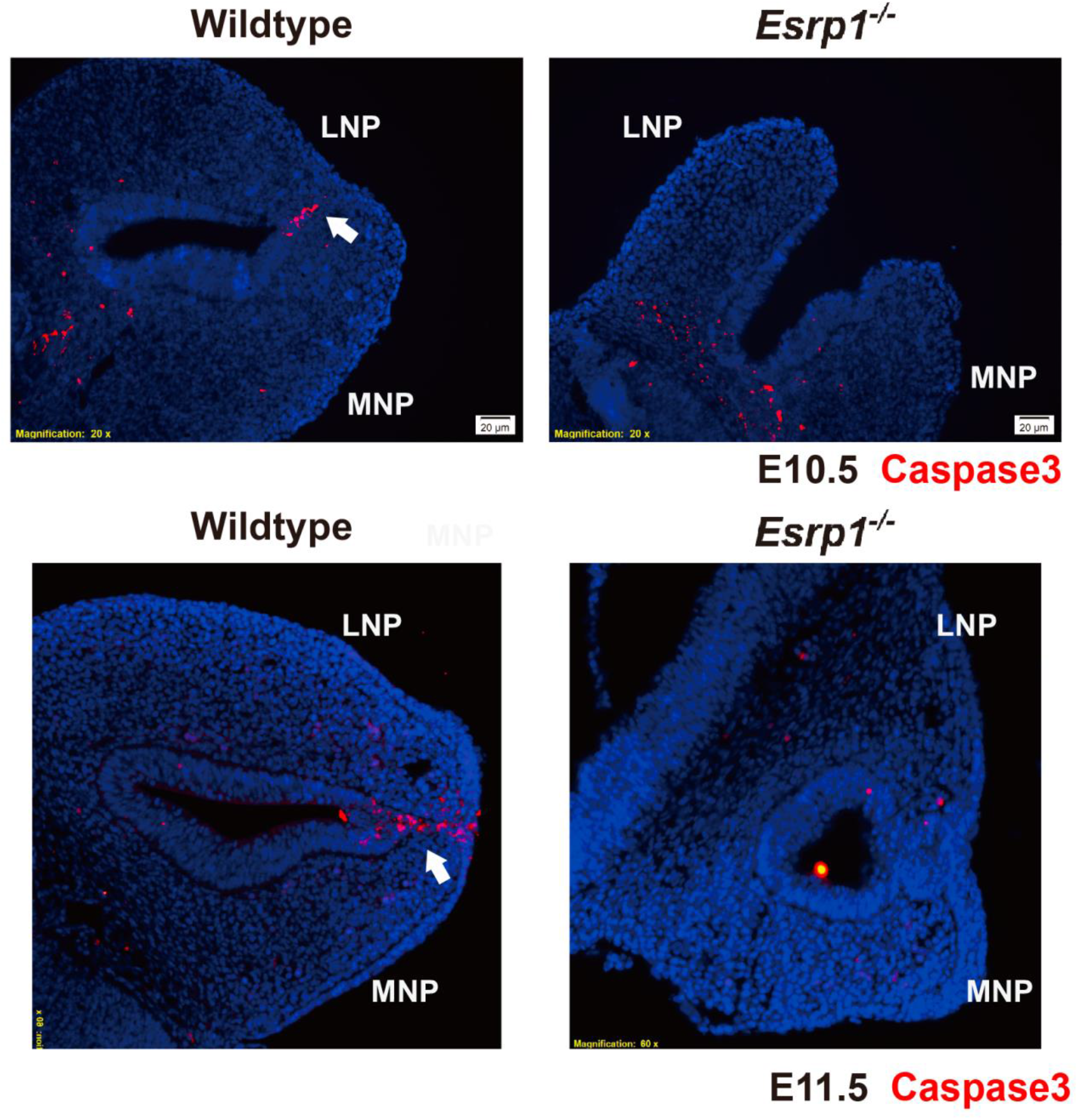
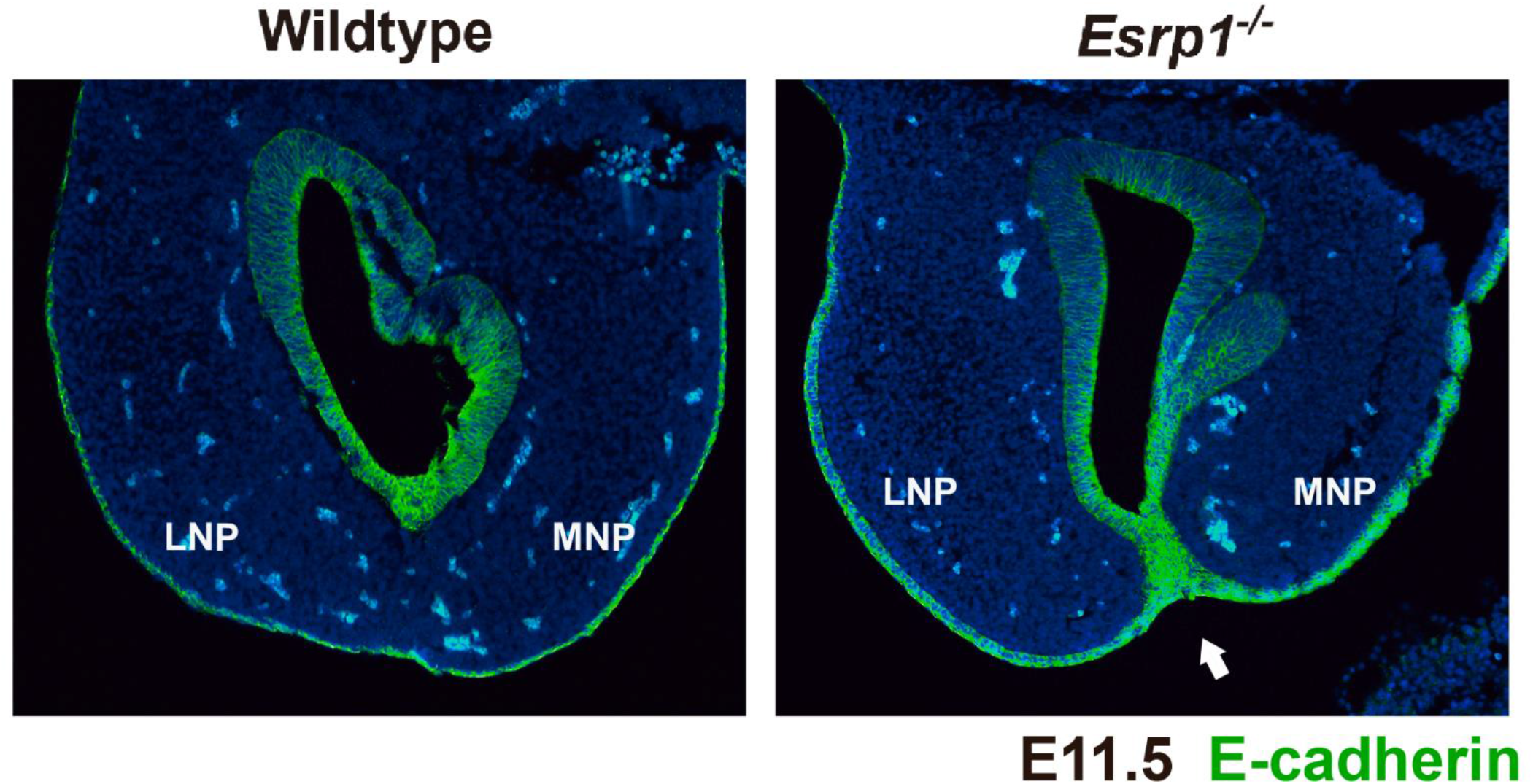
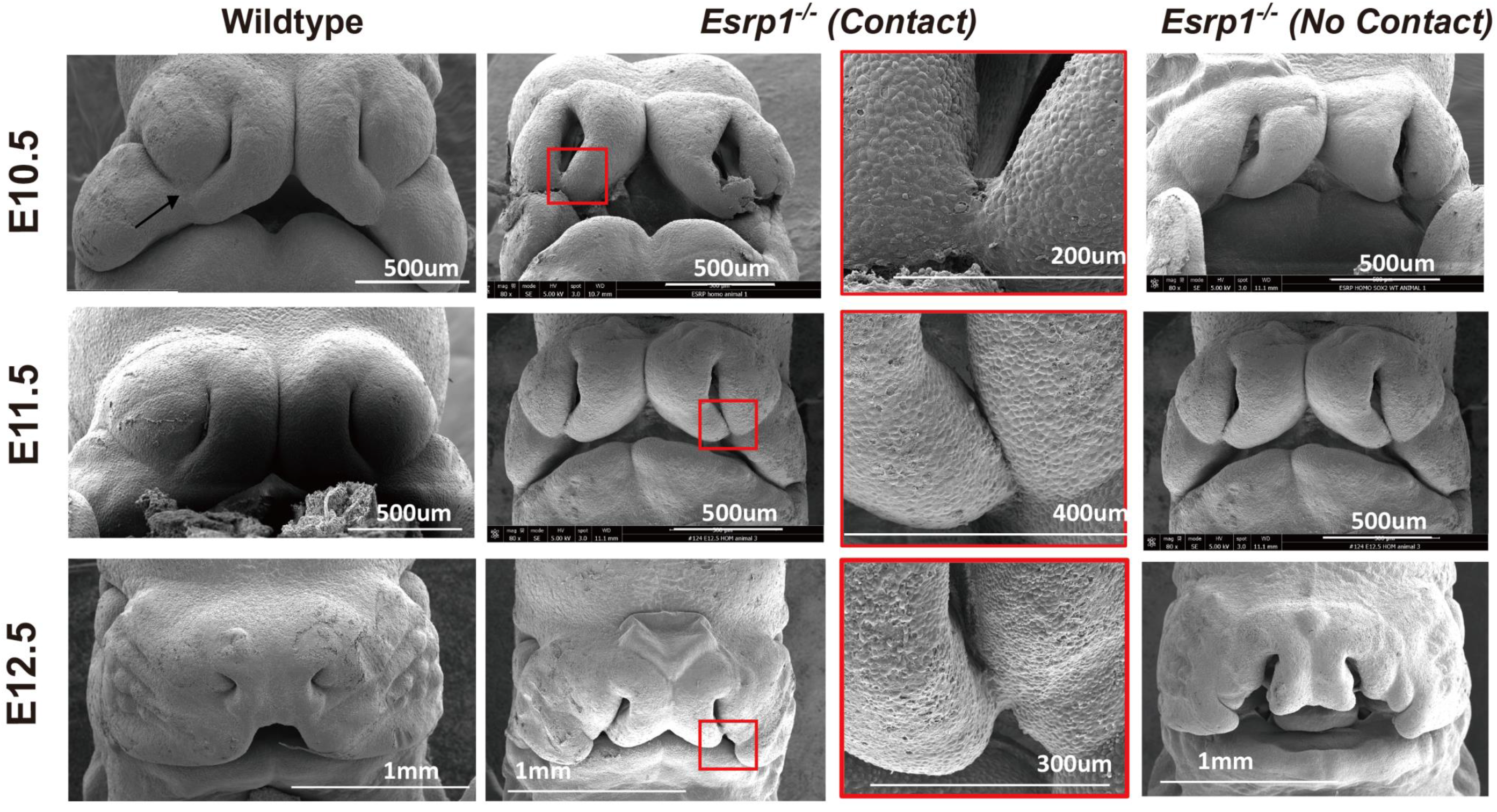
*Esrp1*^−/−^ embryos exhibit reduced proliferation of the MNP and LNP and are unable to fuse despite some contact. (A) Frontal section showing decreased Ki-67 staining (green) in *Esrp1*^−/−^ ectoderm and mesenchyme of *Esrp1*^−/−^ embryos at E10.5 compared to controls. At bottom are quantifications of Ki-67 signal in ectoderm and mesenchyme from three independent WT and *Esrp1*^−/−^ embryos. Y-axis indicates the % of Ki67 positive cells compared to total cells within each section. Error bars indicate standard deviation. Statistical significance comparing each Esrp1−/− sample with wild-type control was determined by t-test. *P <0.05. (B) Arrow shows epithelial seam at fusion site between MNP and LNP in two WT embryos showing with the expected apoptotic cells by staining for activated Caspase 3 (red). MNP and LNP from two *Esrp1*^−/−^ embryos did not reveal substantial apoptosis, including one example where the MNP and LNP make contact. Nuclei are stained with DAPI (blue) (C) E-Cadherin staining (green) showing loss of epithelial cells between MNP and LNP after fusion in WT embryos at E11.5, but persistence of E-cadherin positive epithelial cells in MNP and LNP that made contact *Esrp1*^−/−^ embryos. Nuclei are stained with DAPI (blue) (D) Scanning electron microscopy (SEM) showing normal lip fusion in WT embryos from E10.5 to E12.5. The arrow in the WT image at E10.5 shows the normal formation of the lambdoidal junction. In *Esrp1*^−/−^ embryos we observed examples with no apparent contact between MNP and LNP or cases of some contact, but none with fusion by E12.5. LNP, lateral nasal process; MNP, medial nasal process.

We used scanning electron microscopy (SEM) to further evaluate lip formation and fusion in WT and *Esrp1*^−/−^ embryos at different stages of lip formation. At E10.5, WT embryos showed the onset of fusion between the MNP and LNP, LNP and MxP, and MNP and MxP at the 3-way lambdoid junction. At E11.5 the fusion between these processes was more complete and by E12.5 there was normal development of the upper lip and nasal pit (Fig. 2D). In *Esrp1*^−/−^ embryos the LNP and MNP were smaller and there remained a gap between them that resulted in a larger nasal pit (Fig. 2D). Compared to WT embryos, the MNP and LNP were further separated from each other, but in some cases there was contact between these processes, but no apparent fusion. There was also no contact observed between the hypoplastic MxP and either MNP or LNP to generate a typical lambdoid junction. By E12.5 there was very little contact noted between any of the prominences other than one example where there was some contact between MNP and LNP deeper into the nasal pit. Taken together, the SEM studies showed that while there was some reduction in size of the facial prominences that reduced contact, there were some cases where contact did occur between MNP and LNP, but this was not followed by fusion. These observations, together with sections of these prominences in E10.5 embryos, suggested that while reduced proliferation is likely to partially underlie the CL/P phenotype, there also was an apparent defect in the ability of the epithelial cells of the apposed facial processes to fuse to form the lip.

### Palatal processes show reduced proliferation as well as a defect in fusion during palatogenesis

The studies using SEM and histology suggested that the CL/P observed in *Esrp1*^−/−^ mice was due to both a defect in fusion and in cell proliferation. We also noted reduced outgrowth of the palatal shelves towards each other, indicating that they also likely had a defect in cell proliferation. Further analysis of the defect in secondary palate formation in *Esrp1*^− −^E16.5 embryos using Ki67 staining and cleaved caspase 3 detection confirmed that the palatal shelves had a proliferation defect without apparent differences in apoptosis compared to wild-type embryos (Fig 3A). However, because the palatal shelves did not make contact *in vivo* in *Esrp1*^−/−^ mice, we were unable to directly determine whether there was an associated fusion defect. Therefore, to further evaluate the defect in palatogenesis in *Esrp1* KO mice we performed palatal organ culture assays. WT and *Esrp1*^−/−^ palatal shelves were isolated from E13.5 embryos and cultured for up to 72 hours. Palatal shelves from WT embryos grew together within 48 hours and after adhering underwent fusion with dissolution of the medial epithelial seam (MES) and mesenchymal confluence (N=7/7). Evaluation of palatal cultures from *Esrp1*^−/−^ mice was complicated by increased fragility and reduced proliferation such that contact between opposing palatal shelves was delayed. Nonetheless, we noted that while the palatal shelves from eight out of thirteen *Esrp1*^−/−^ embryos were able to achieve contact, in seven of the eight palatal cultures where the palatal processes achieved adherence there was no dissolution of the medial edge epithelial cells and a failure in achieving mesenchymal confluence. Only one of the eight *Esrp1*^−/−^ palatal shelves that made contact showed partial dissolution of the MES and a small region of apparent mesenchymal connection. For one representative control and one *Esrp1* KO sample in which the palatal processes made contact, we performed staining for the epithelial marker E-cadherin as well as activated caspase 3 and Ki67. In the wild-type cultures, we observed a nearly complete loss of E-cadherin expression in cells at the site of fusion as well as the expected apoptosis at the fusion site. However, in the *Esrp1* KO palates that achieved close contact, there was persistence of the MES and an absence of mesenchymal confluence (Fig. 3B, additional examples in Fig. S2). These results demonstrate that while reduced growth and proliferation of palatal shelves contribute to the cleft palate defect, there is also a defect in fusion and dissolution of the MES.

**Fig. 3.**
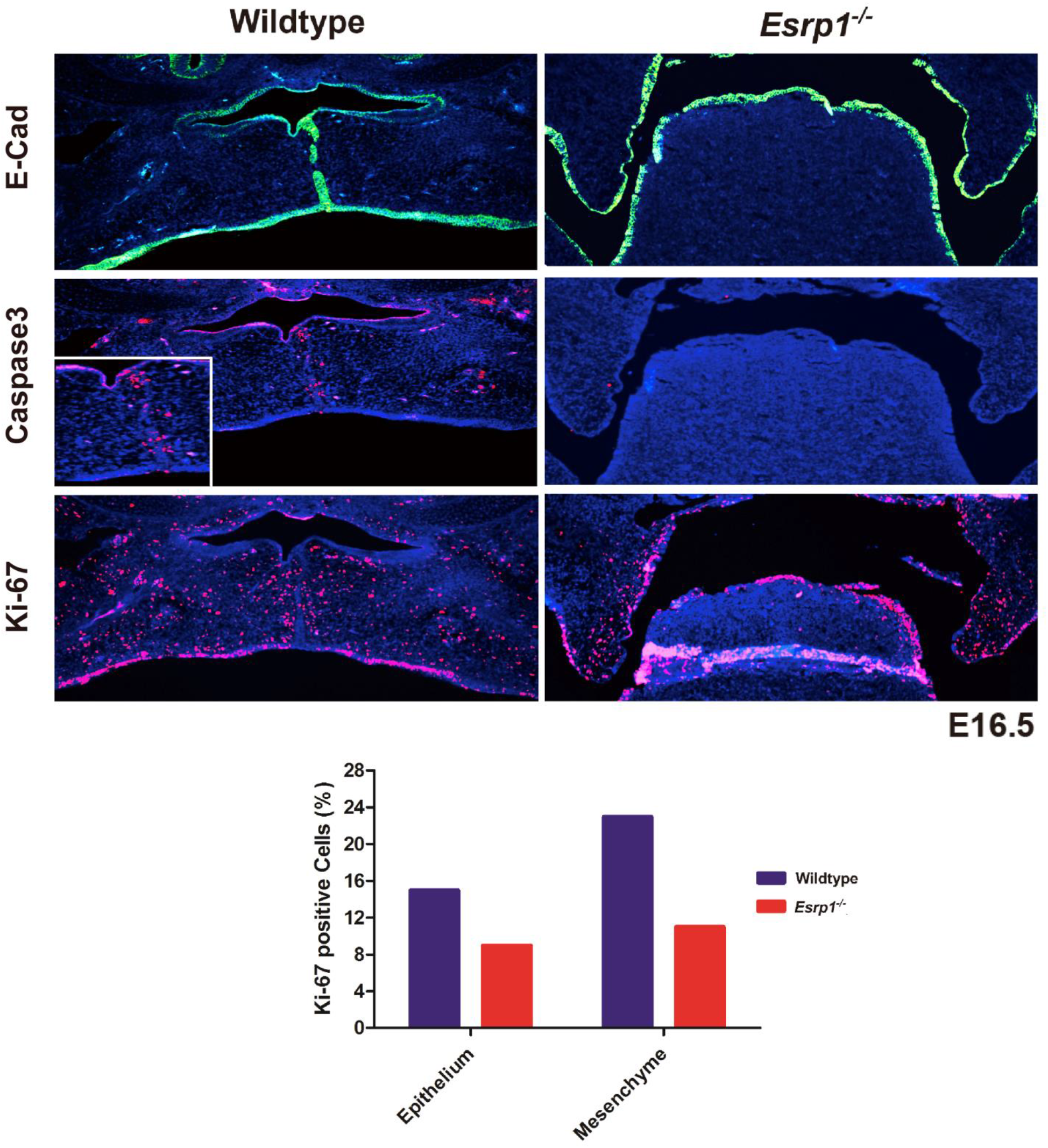
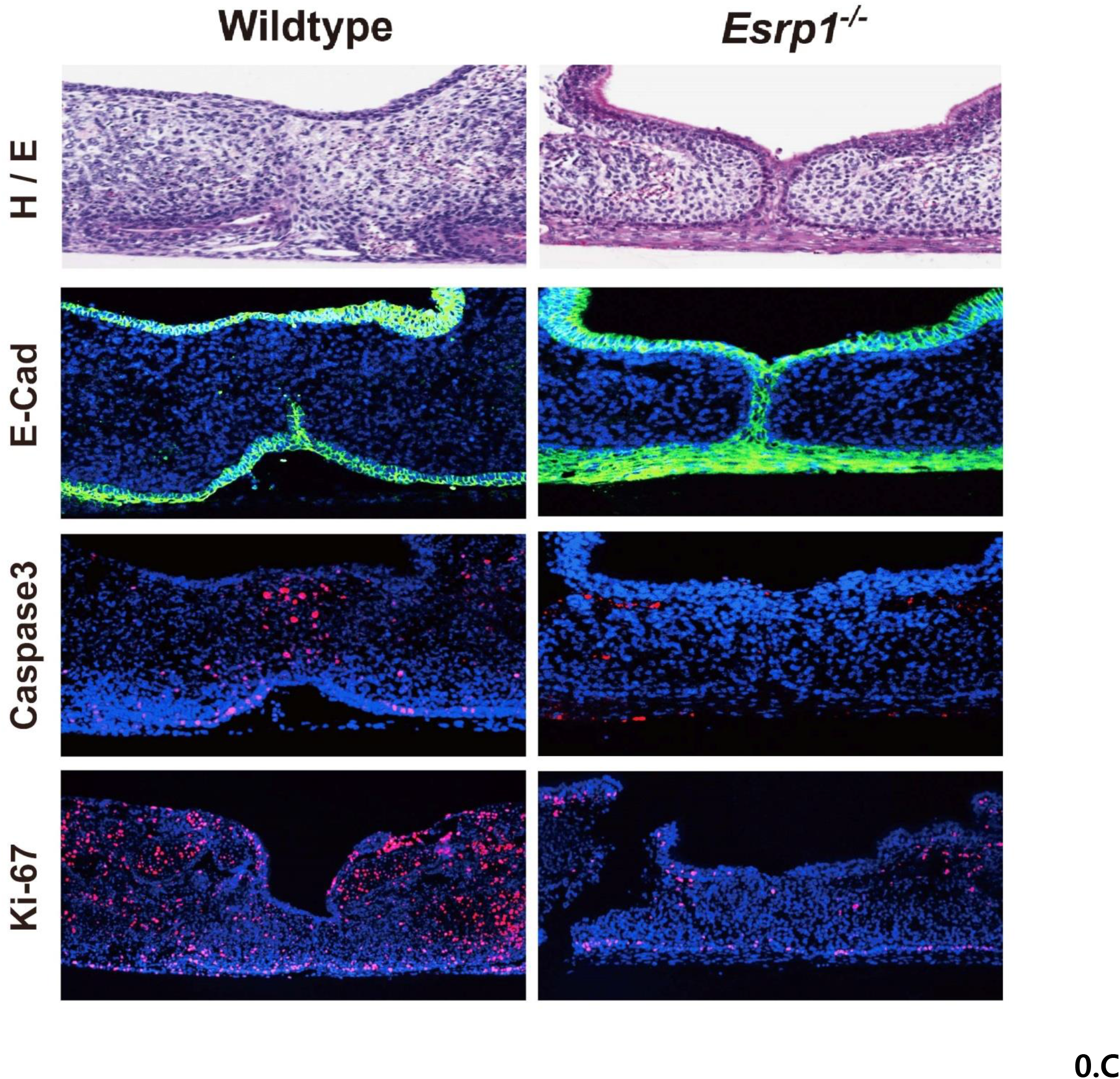
Reduced proliferation and a fusion defect contribute to cleft palate in *Esrp1*^−/−^ mice. (A) Coronal sections with immunofluorescent staining for E-cadherin (green) in E16.5 control and Esrp1−/− embryos indicates the medial edge epithelial seam (MES) is becoming discontinuous following fusion in controls whereas the palatal shelves have failed to elevate in the mutants. Staining for activated Caspase 3 shows apoptosis at the site of fusion, but no apparent increase in apoptosis in *Esrp1*^−/−^ embryos, which also show reduced Ki-67 staining for proliferating cells compared to WT in the palatal shelves. Nuclei are stained with DAPI (blue). (B) Palatal organ culture showing lack of dissolution of the MES and associated apoptosis and reduced proliferation in palatal shelves from *Esrp1*^−/−^ embryos compared to WT.

### Ablation of Esrp1 leads to large scale changes in splicing in surface ectoderm

The specific expression of ESRP1 in surface ectoderm together with the CL/P defect observed in *Esrp1*^*flox/flox*^;*Crect*^+/−^ embryos indicated that alterations in the transcriptome of surface ectoderm derived cells underlie orofacial clefting in *Esrp1*^−/−^ mice. However, the reduction in cell proliferation observed in the mesenchyme adjacent to epithelial cells also indicated that gene expression alterations in *Esrp1*^−/−^ epithelial cells induced changes in signaling crosstalk from epithelium to mesenchyme. To further investigate the molecular mechanisms that lead to CL/P in *Esrp1*^−/−^ mice, we performed RNA-Seq using RNAs collected from both epithelial cells and mesenchymal cells from control and *Esrp1*^−/−^ embryos. We used a previously described method to separate facial ectoderm and mesenchyme from facial prominences at E12.0, a stage at which lip fusion is underway (Li and Williams, 2013). We collected pooled paired ectoderm and mesenchyme samples to obtain sufficient material for four replicates each of ectoderm and mesenchyme fractions from WT and *Esrp1*^−/−^ embryos and prepared total RNA for RNA-Seq. We used paired end sequencing and obtained deep coverage with an average of 100 million read pairs per replicate. Preliminary analysis of transcripts per million (TPM) values in the RNA-Seq analysis from epithelial and mesenchymal control samples validated that they were derived from relatively pure populations of each cell type using a panel of standard epithelial and mesenchymal cell type-specific markers, including *Esrp1* (Fig. 4A). To identify genome-wide alterations in splicing in ectoderm from *Esrp1* KO embryos compared to WT controls we used replicate-based multivariate analysis of transcript splicing (rMATS)(Shen et al., 2014). We also used rMATS to identify global differences in splicing of epithelial cells compared to mesenchymal samples using RNAs from control embryos. In *Esrp1* KO epithelial cells rMATS identified a total of 1467 alternative splicing changes compared to wild-type (WT) epithelial cells, with cassette exons (skipped exons (SE)) representing the largest fraction (Fig.4B, 4C, Table S1). Analysis of splicing differences between WT epithelial cells and WT mesenchymal cells identified 2546 splicing events, including many of the Esrp-regulated events that switch splicing from the epithelial to mesenchymal splice variants after Esrp1 ablation (Fig. 4B, 4D, Table S2). The larger number of splicing differences between epithelial to mesenchymal cells compared to *Esrp1*^−/−^ epithelial cells is consistent with our previous studies showing larger changes in splicing during the epithelial to mesenchymal transition (EMT) than when *ESRP1* and *ESRP2* were depleted in the same cell line (Yang et al., 2016). These observations are also in line with previous studies by our group and other investigators showing combinatorial regulation of splicing events that are induced during EMT or that differ between epithelial and mesenchymal cells. This combinatorial regulation includes the ESRPs as well as other splicing factors such as RBFOX2, QKI, RBM47, and MBNL1 (Braeutigam et al., 2014; Pillman et al., 2018; Shapiro et al., 2011; Venables et al., 2013; Yang et al., 2016). We validated 17 cassette exon (SE) events in WT compared to *Esrp1*^−/−^ ectoderm along with 6 examples of splicing differences between normal ectoderm vs. normal mesenchyme by semi-quantitative RT-PCR (Fig. 4E,F). As expected, a change in alternative splicing of *Fgfr2* was identified among alterations in mutually exclusive (MXE) exon splicing in *Esrp1*^−/−^ ectoderm. This event involves two alternative exons, named IIIb or IIIc, that encode a region in the extracellular ligand binding domain and with the resulting receptor isoforms, FGFR2-IIIb and FGFR2-IIIc, having different FGF binding preferences (Zhang et al., 2006). We also confirmed this splicing switch by RT-PCR, which showed a switch from nearly complete splicing of the epithelial-specific IIIb exon to predominant splicing of the mesenchymal IIIc in *Esrp1*−/− ectoderm (Fig. 4E). Both global splicing comparisons were then subjected to Gene Ontology analysis and revealed overlapping enrichment terms for both splicing comparisons, including cytoskeletal organization, cell morphogenesis, and cellular component organization (Fig. 4G, H). Similarly, Kyoto Encyclopedia of Genes and Genomes (KEGG) pathway analysis demonstrated an enrichment for terms including adherens junctions, suggesting that alternatively spliced genes contributed to epithelial functions, including regulation and/or assembly of this structure (Fig. 4G, H).

**Fig. 4.**
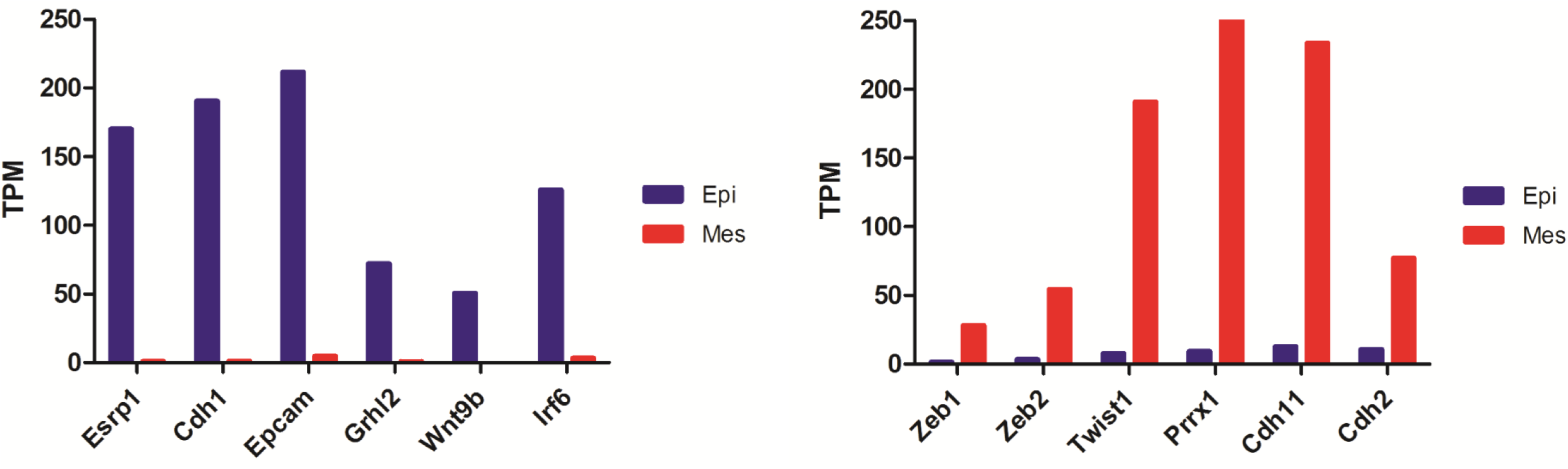
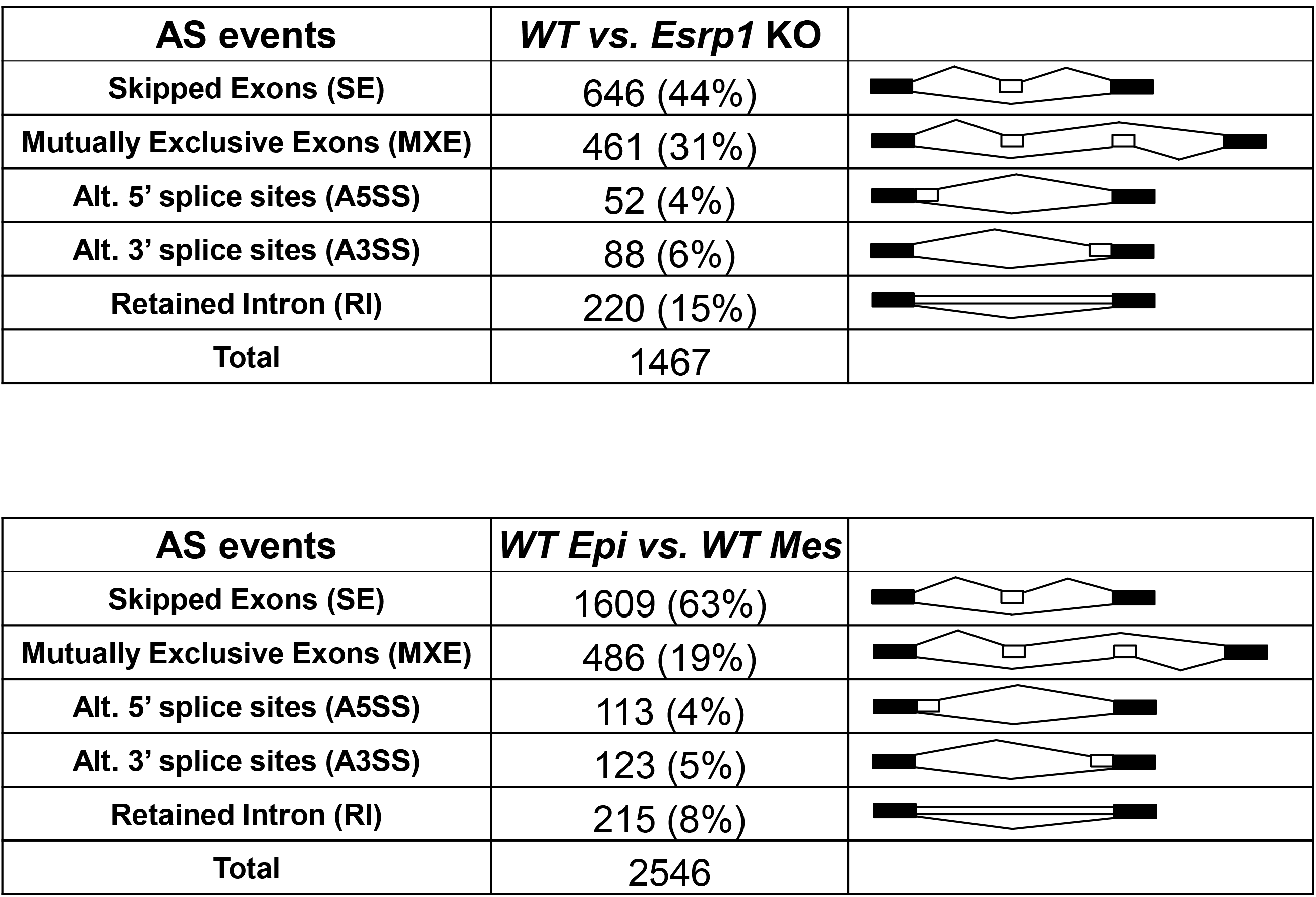
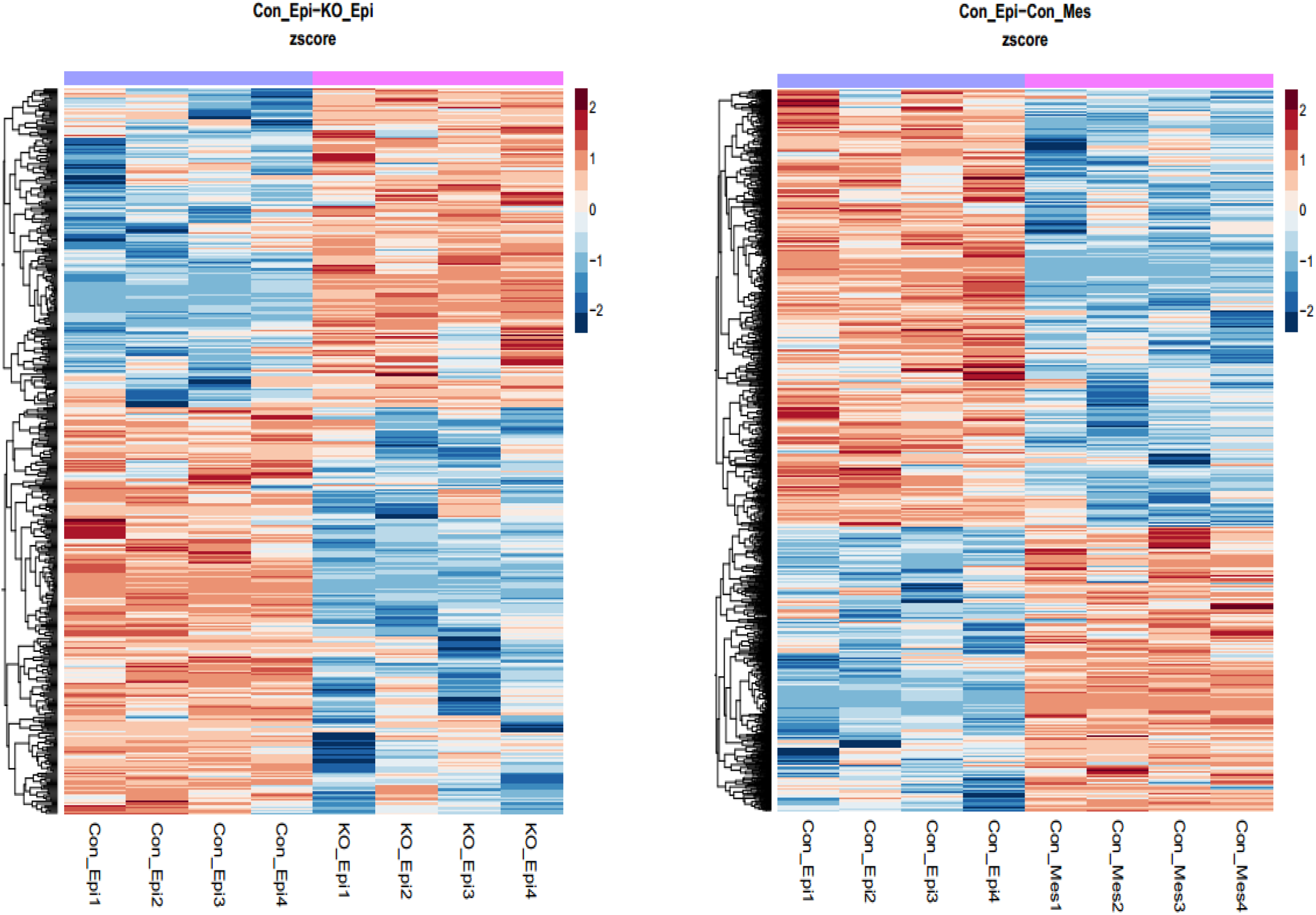
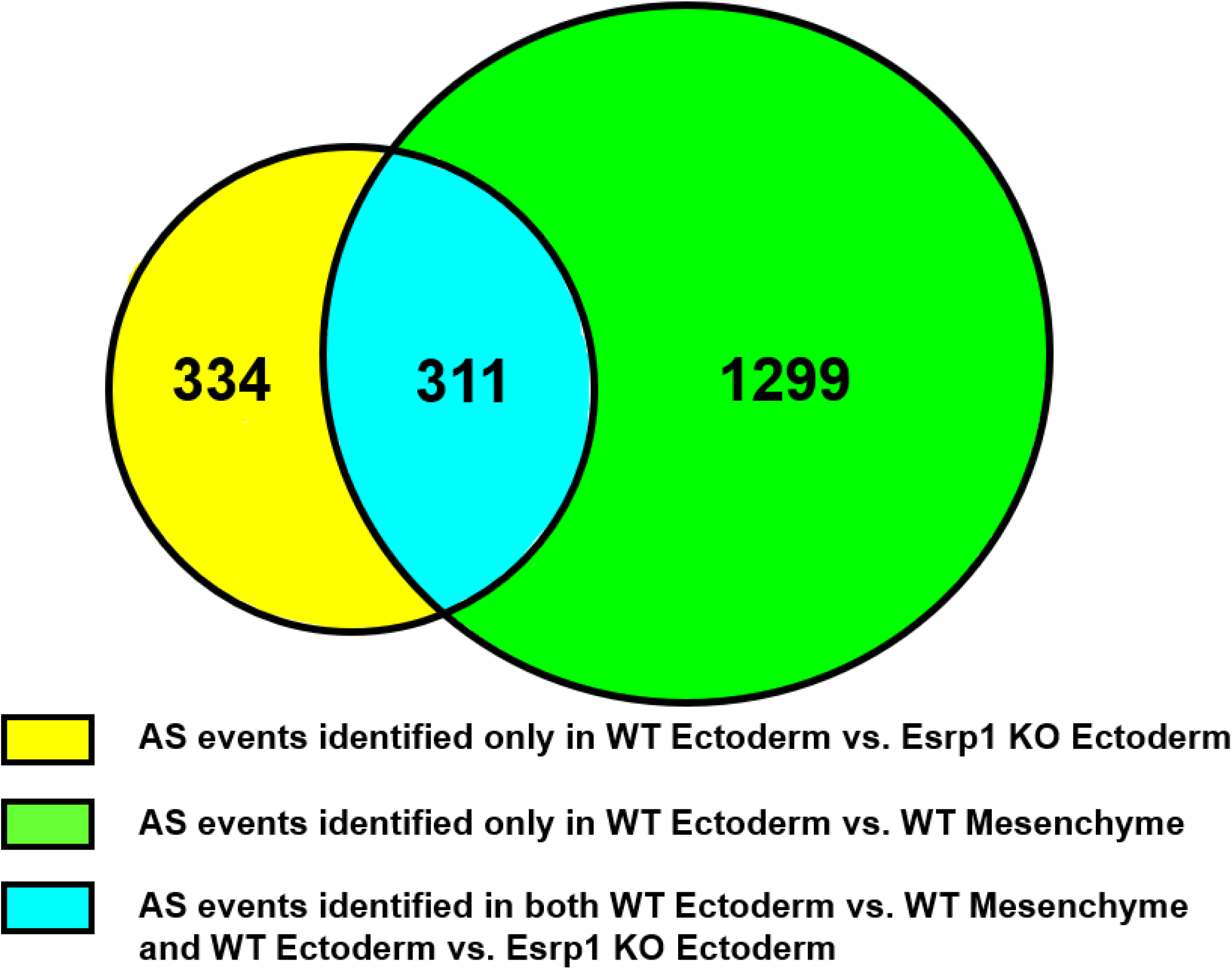
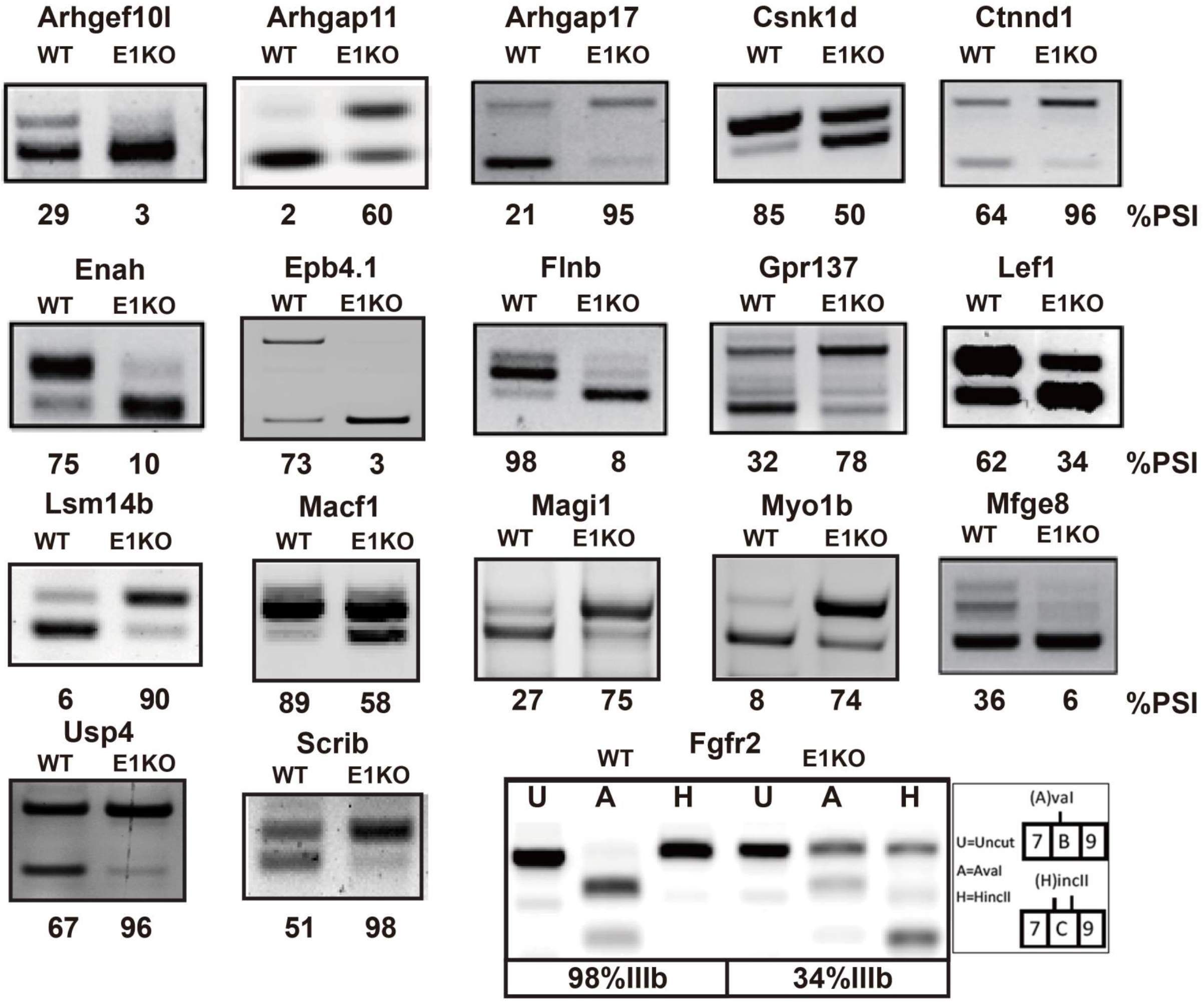
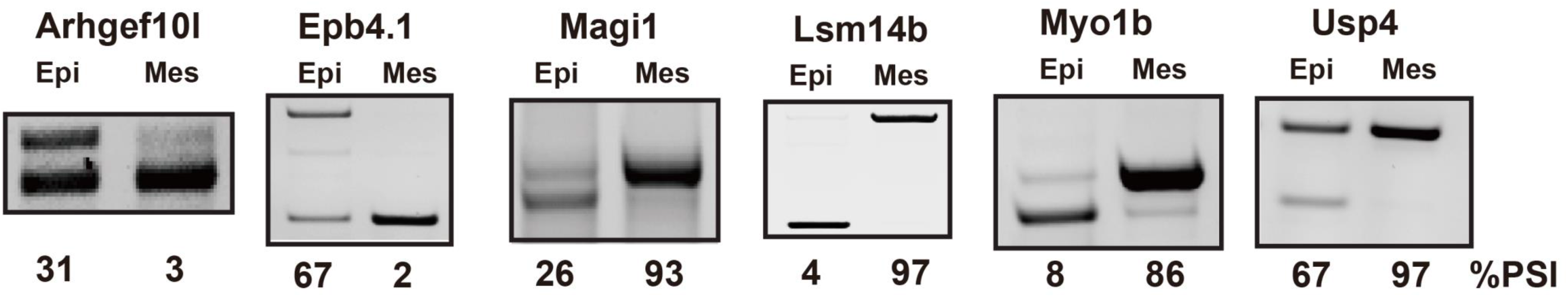
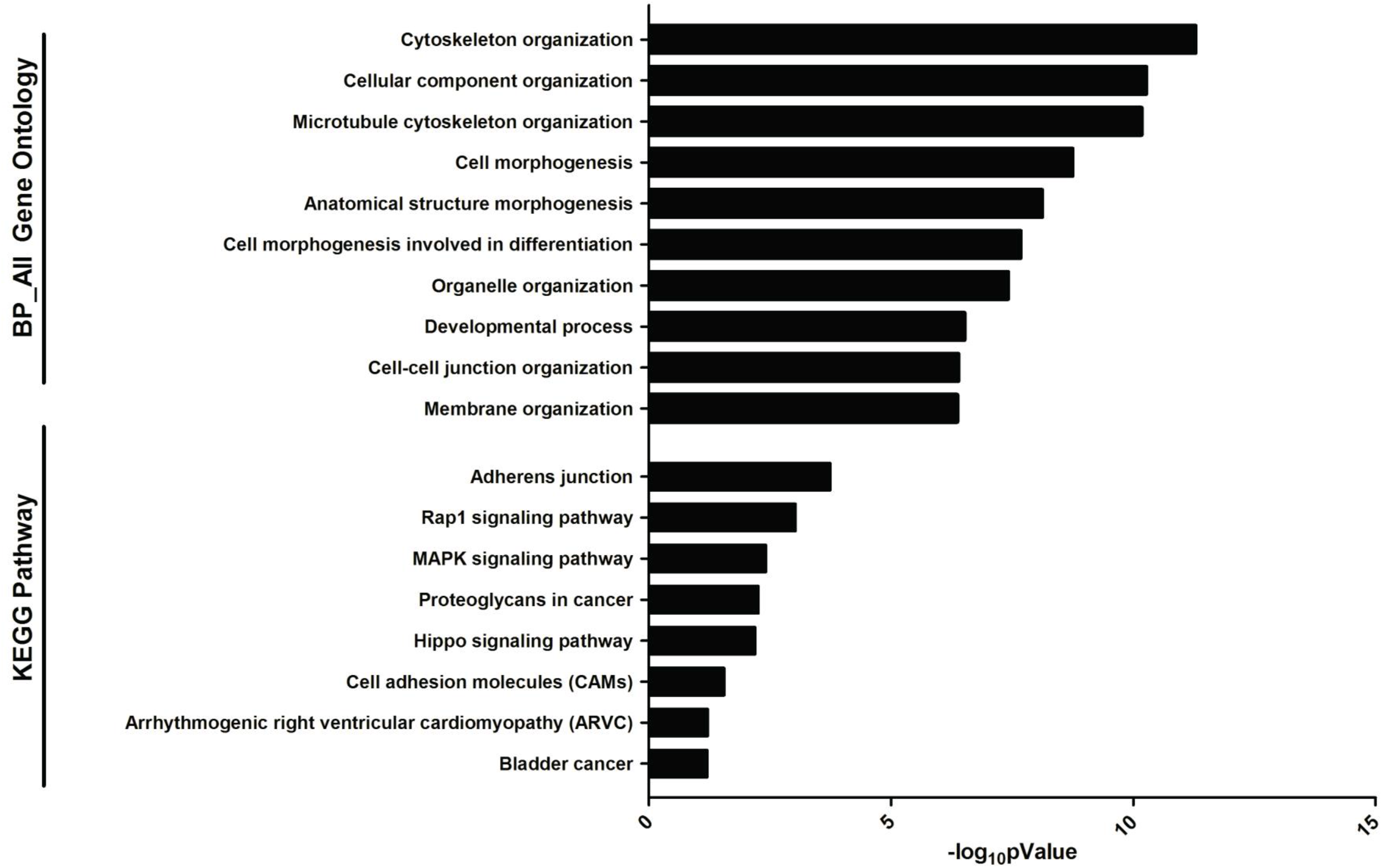
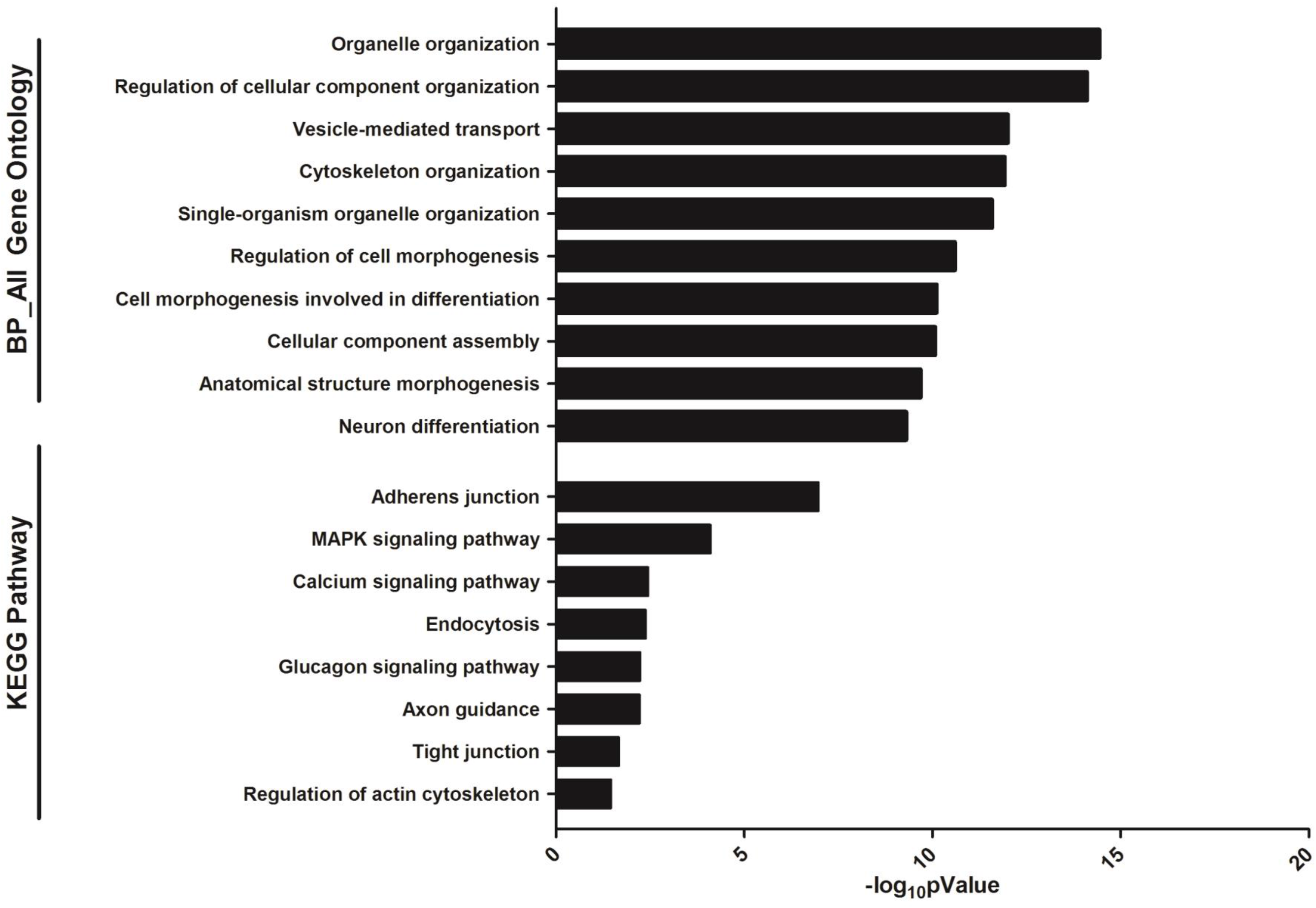
RNA-Seq from epithelial and mesenchymal cells identified large scale alterations in alternative splicing in *Esrp1*^−/−^ ectoderm compared to WT ectoderm and differences in alternative splicing between WT ectoderm and WT mesenchyme. (A) Expression of a panel of epithelial specific markers (left) and mesenchymal markers (right) validates efficient separation of epithelial cells from mesenchymal cells in WT samples. TPM = transcripts per million. (B) Summary table of different types of alternative splicing (AS) events in WT vs. *Esrp1*^−/−^ ectoderm and in WT ectoderm vs. WT mesenchyme. Results obtained using rMATS with False Discovery Rate (FDR) <5%, and |deltaPSI|>=5%. (C) Heatmap representing the skipped exon (SE) splicing changes with increased exon inclusion (red) or decreased exon inclusion (blue). (D) Venn diagram depicting detected SE events identified in WT ectoderm vs. Esrp1 KO ectoderm, WT ectoderm vs. WT mesenchyme, and those identified in both comparison sets. (E) RT-PCR validations of changes in splicing of cassette exons *Esrp1*^−/−^ ectoderm (E1KO) compared to WT. The Percent Spliced In (PSI) value indicates % exon inclusion for each event. Also shown is the change in splicing of mutually exclusive exons IIIb and IIIc of Fgfr2 from predominant use of the IIIb exon, to mostly exon IIIc splicing. Products containing each exon were distinguished by restriction digests with Ava I and Hinc II, which cut products containing exon IIIb and IIIc, respectively. (F) Validation of several differences in splicing between control ectoderm and mesenchyme. (G) Gene Ontology (GO) and KEGG pathway enrichment for genes with alternative splicing differences between WT ectoderm and Esrp1^−/−^ ectoderm. H. Gene Ontology (GO) and KEGG pathway enrichment for genes with splicing differences between WT ectoderm and WT mesenchyme.

### Ablation of Esrp1 in ectoderm reduces expression of canonical Wnts and is associated with reductions in canonical Wnt target transcripts in underlying mesenchyme

We also investigated changes in total transcript levels in *Esrp1* ablated epithelial cells, as well as in adjacent mesenchyme. In epithelial cells we identified 167 downregulated genes and 546 upregulated genes (Table S3). Among the downregulated genes we noted six transcripts encoding canonical Wnt ligands, Wnt3, Wnt3a, Wnt4, Wnt7b, Wnt9b, and Wnt10b, each of which were downregulated approximately 2-fold, of which several were validated by RT-qPCR (Fig. 5A). Furthermore, we also noted downregulation of Wls, which is required for Wnt secretion. Consistent with these observations, for genes downregulated in *Esrp1* KO ectoderm, Wnt signaling pathway was the most enriched gene ontology (GO) term for biological process and was also enriched by KEGG pathway analysis (Fig. 5B). We also noted an increase in both *Aldh1a2* and *Aldh1a3*, which generate retinoic acid and have been implicated in feedback regulation of Wnt signaling during craniofacial development (Osei-Sarfo and Gudas, 2014; Song et al., 2009). We further examined whether the downregulation of Wnt ligands in the ectoderm was also associated with a reduction in targets of canonical Wnt/beta-catenin signaling in mesenchyme as previously shown in *Wnt9b* and *Lrp6* KO mice with CL/P (Jin et al., 2012). Interestingly, we identified larger changes in gene expression in mesenchyme of *Esrp1 KO* mice than in ectoderm, with 3048 upregulated genes and 2396 downregulated genes using the same filtering criteria (Table S3). In addition we also identified genes that were differentially expressed in wild-type ectoderm compared to wild-type mesenchyme, which included numerous known epithelia and mesenchymal markers in addition to those indicated in Fig. 4A (Table S5). While Wnt signaling was not among the most enriched GO terms or pathways for genes downregulated in *Esrp1*^−/−^ mesenchyme compared to wild-type mesenchyme, we noted numerous canonical Wnt targets among downregulated genes under the “Wnt signaling pathway” category, several of which were also validated by RT-qPCR along with several other genes encoding components of the Wnt pathway (Fig. 5A). We also noted that there was reduced expression of Sonic hedgehog (Shh) in *Esrp1* KO ectoderm and a corresponding reduction in expression of the *Gli1*, *Gli2* and *Gli3* transcription factors in mesenchyme, with the greatest reduction in *Gli2* confirmed by RT-qPCR (Fig. 5A). Previous studies have identified crosstalk between Wnt and hedgehog signaling and ablation of Shh in palatal epithelial cells or in utero treatment with Shh inhibitors, have been shown to cause cleft palate and CL/P, respectively (Lan and Jiang, 2009; Lipinski et al., 2010; Reynolds et al., 2019; Rice et al., 2004). It is thus possible that reduction in Shh expression in ectoderm might also contribute to CL/P in *Esrp* KO mice. We also identified several other genes with reduced expression in *Esrp1* KO mesenchyme that have previously been implicated in cleft palate, including *Bmp7*, *Tgfb2*, and *Tgfb3* (Fig. 5A, Table S4) (Kaartinen et al., 1995; Kouskoura et al., 2013; Proetzel et al., 1995; Sanford et al., 1997). Thus, while reductions in components of the canonical Wnt signaling pathway in ectoderm were the most conspicuous pathway that has been strongly linked to CL/P, there were also alterations in a number of other pathways and genes implicated in cleft lip and/or palate. A future challenge will be to dissect the potential contributions of different pathways whose disruption in *Esrp1* KO mice plays a role in CL/P.

**Fig. 5.**
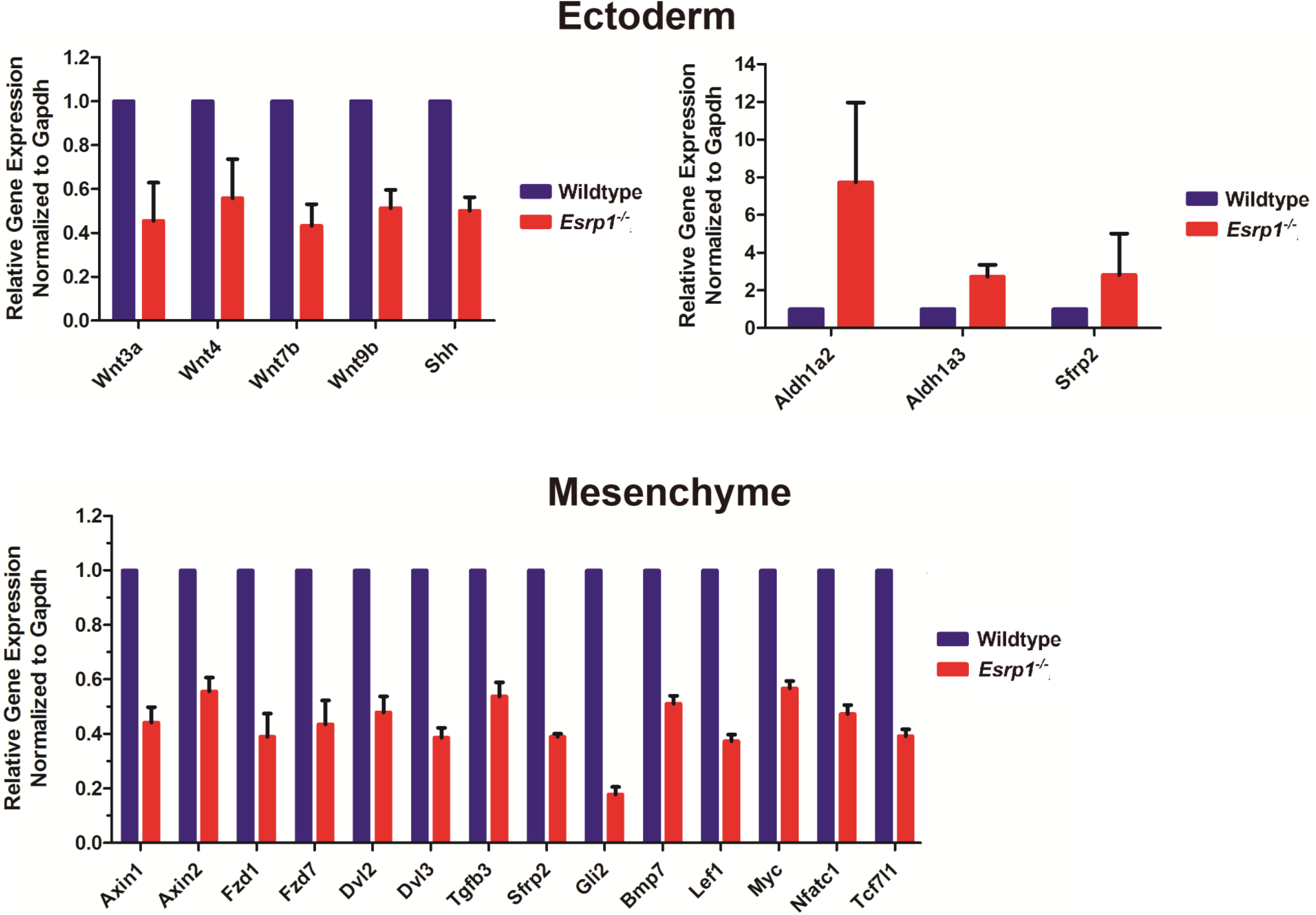
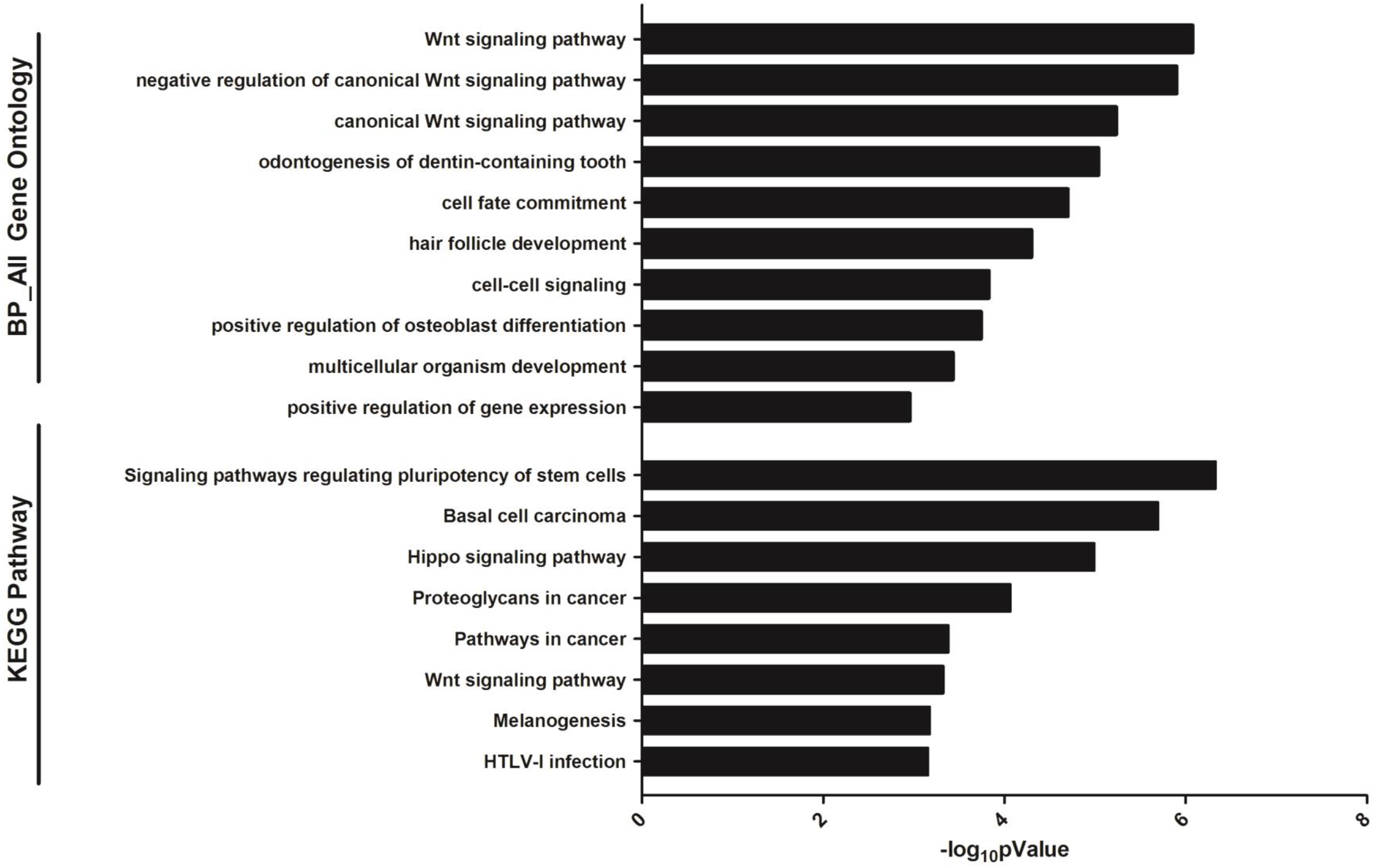
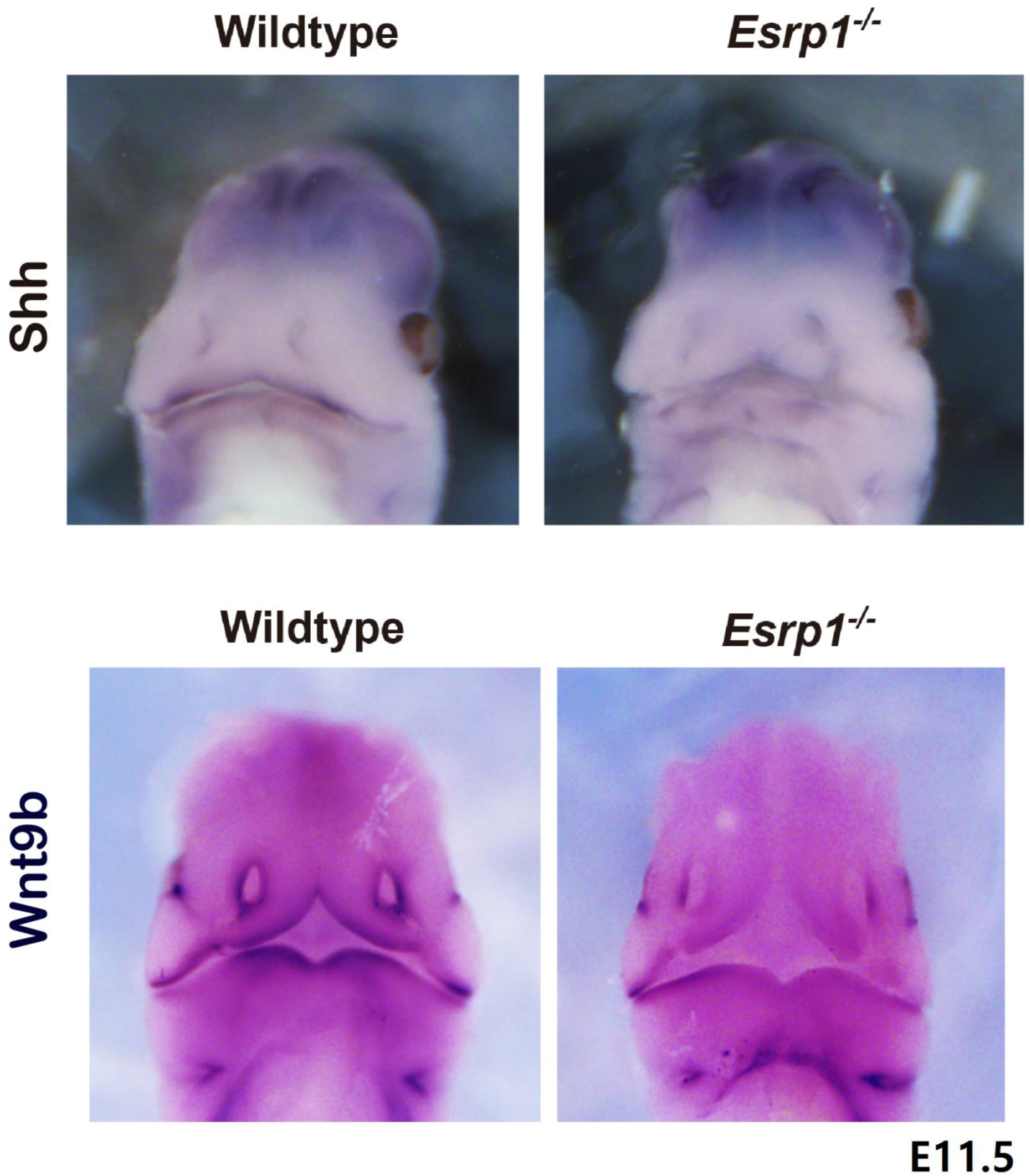
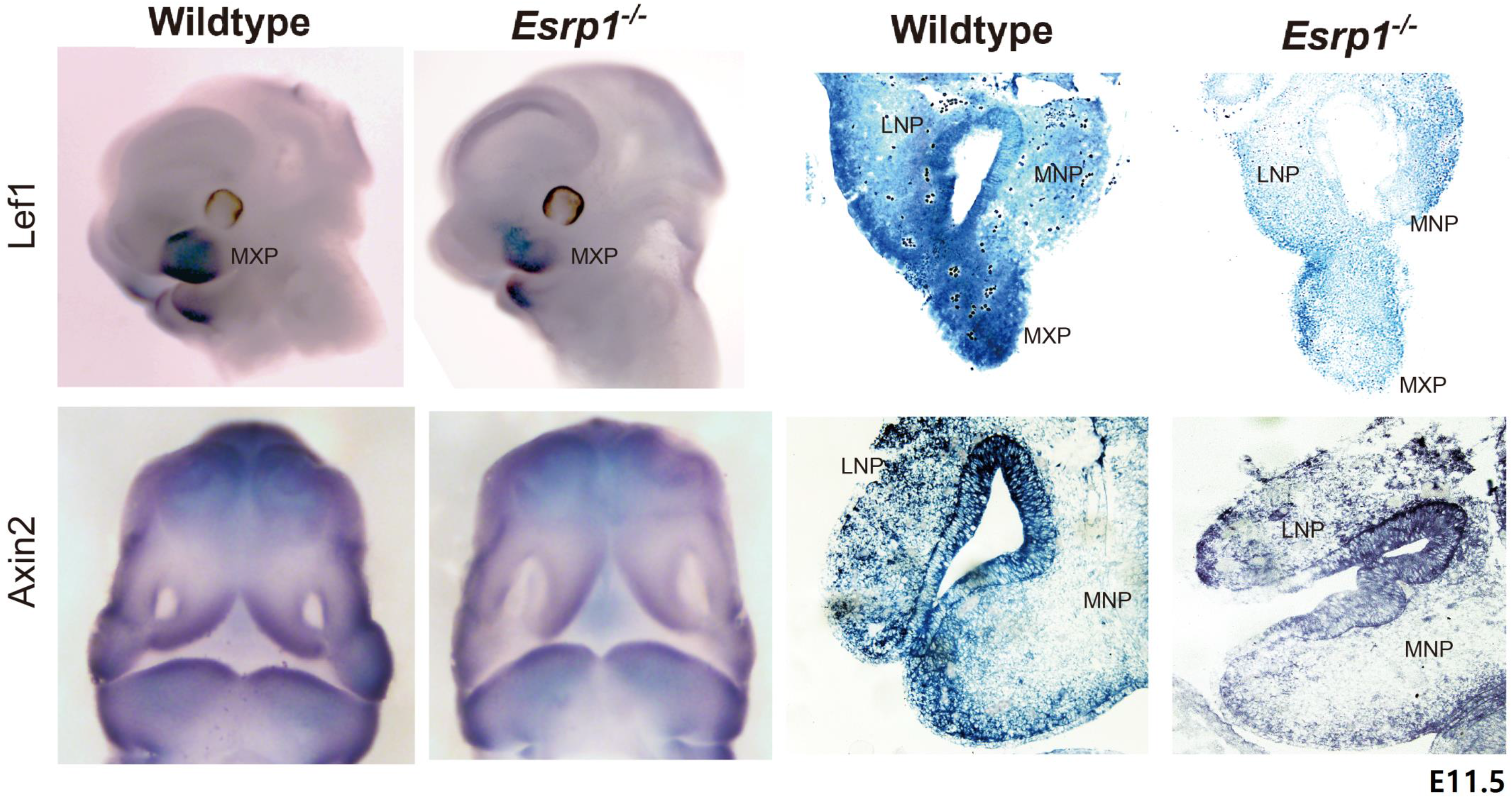
Esrp1 ablation in ectoderm results in altered expression of components of the Wnt signaling pathway as well as other signaling pathways implicated in cleft lip and/or cleft palate. (a) qRT-PCR validations for select changes in total transcript levels in *Esrp1*^−/−^ ectoderm as well as in adjacent mesenchyme. Error bars indicate standard deviation. (B) Gene Ontology (GO) and KEGG pathway enrichment for genes that are downregulated in *Esrp1*^−/−^ ectoderm. (C) Whole mount in situ hybridization of E11.5 embryos showing reduced expression of Shh and Wnt9b in epithelial cells of the developing face. (D) Whole mount and tissue sections showing reduced expression of canonical Wnt target genes *Lef1* and *Axin2* in *Esrp1*^−/−^ embryos. LNP, lateral nasal process; MNP, medial nasal process, MxP, maxillary process.

We validated reduced expression of the Wnt signaling pathway as well as Shh in *Esrp1*^−/−^ mice using in situ hybridization (ISH) in mouse embryos at E10.5-11.5. Whole mount ISH confirmed a reduction in both *Wnt9b* and *Shh* that was limited to facial ectoderm in *Esrp1*^−/−^ embryos, which was also consistent with the RNA-Seq data showing that both *Wnt9b* and *Shh* were specifically expressed in ectoderm (Fig. 5C, Table S3). We also confirmed reduced expression of canonical Wnt targets *Lef1* and *Axin2* using both whole mount and section ISH (Fig. 5D).

To further examine canonical Wnt signaling during facial development in *Esrp1*^−/−^ mice, we also crossed both WT and *Esrp1*^−/−^ mice with *TCF/Lef:H2B-GFP* transgenic reporter mice that express an H2B-EGFP fusion protein under the control of six copies of the TCF/LEF response element(Ferrer-Vaquer et al., 2010). In *Esrp1*^−/−^ embryos at E11.5, we noted reduced reporter activity in both the ectoderm and mesenchyme of the NPs and MxP, although the reduction in mesenchyme was more pronounced in MxP than the MNP or LNP (Fig. 6A). Because of some differences in various Wnt reporter models and to further verify changes in canonical Wnt signaling, we also used *Axin2lacZ* mice in which LacZ is knocked in at the endogenous *Axin2* locus as a second readout for Wnt signaling in WT vs. *Esrp1*^−/−^ embryos. At E10.5 we noted that although there was no apparent difference in LacZ expression in the LNP, there was pronounced reduction in LacZ in both the ectoderm and mesenchyme of the MNP (Fig. 6B). Furthermore, we also noted reduced LacZ in the MxP at E11.5. Taken together, these observations are consistent with the results from RNA-Seq and show that there is reduced Wnt expression in ectoderm of *Esrp1*^−/−^ embryos and an associated reduction in canonical Wnt targets consistent with a model in which Esrp ablation leads to alterations in in epithelial-mesenchymal interactions that underlie normal face facial development.

**Fig. 6.**
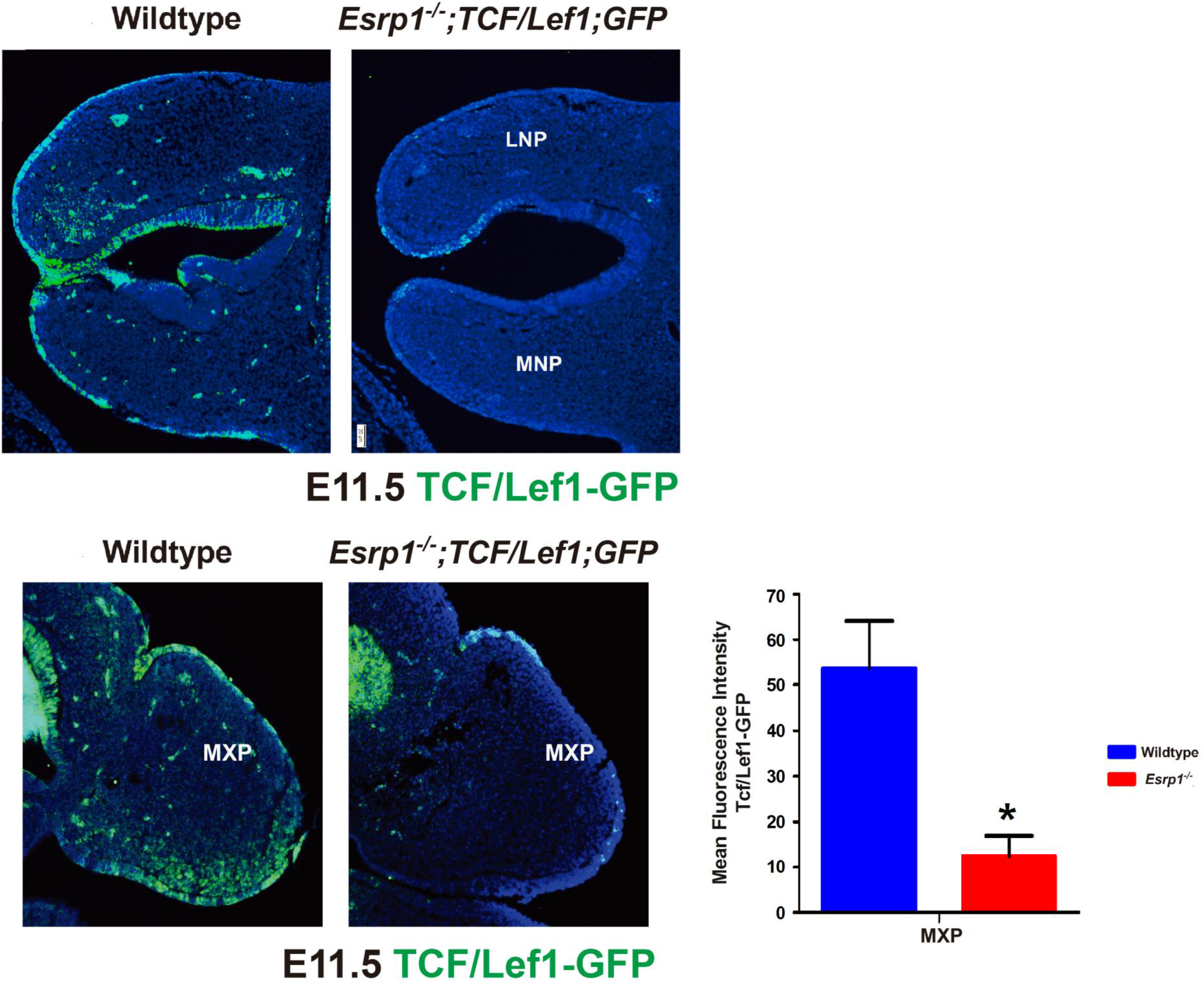
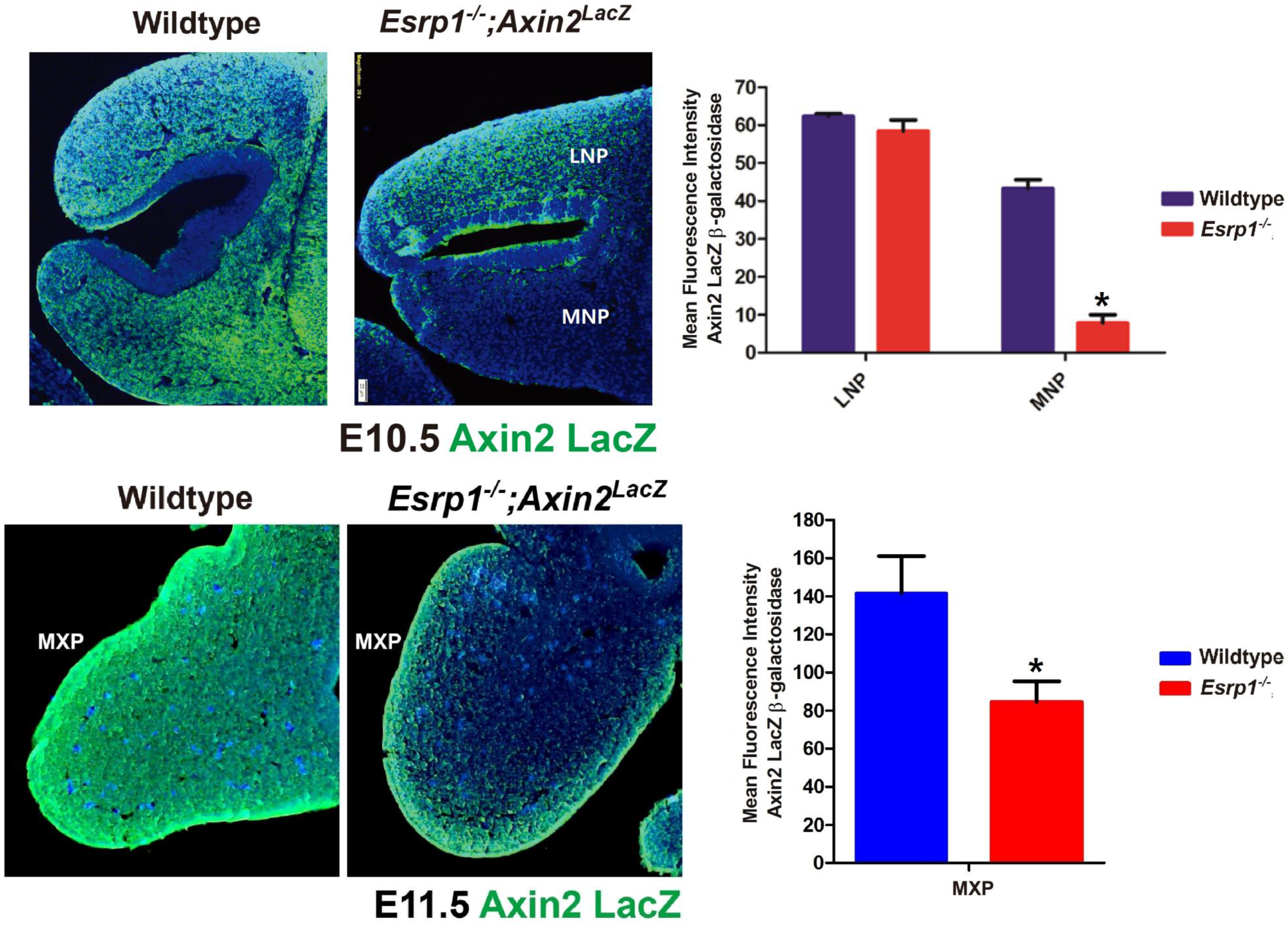
Reduced Wnt signaling in *Esrp1*^−/−^ confirmed in crosses with WNT/β-catenin signaling reporter mice. Frontal sections (A) Reduced activation of the TCF/Lef1-GFP reporter is observed in ectoderm and mesenchyme of *Esrp1*^−/−^ MNP, LNP, and MXP. (B) Reduced expression of the Axin2-LacZ reporter in LNP and MXP in *Esrp1*^−/−^ embryos. Nuclei are stained with DAPI (blue). Error bars indicate standard deviation. Error bars indicate standard deviation. Statistical significance comparing each Esrp1−/− sample with wild-type control was determined by t-test. *P <0.05. Mean Fluorescence Intensity was corrected for area.

## DISCUSSION

Studies using mouse models have played a major role in our understanding of the morphogenetic programs in the face and palate that, when disrupted, lead to cleft lip with or without cleft palate. Mouse models have been particularly informative in defining genes whose ablation leads to cleft palate only (CPO) and collectively these studies have defined molecular pathways that are essential for normal palatal development (Li et al., 2017). However, there remain relatively few mouse models that lead to CL/P, such as those involving mutations or deletions of *Wnt9b, Lrp6, Bmp4, Bmpr1a, Tfap2a*, and *Pbx1/2* (Ferretti et al., 2011; Green et al., 2015; Gritli-Linde, 2012; Jin et al., 2012; Juriloff and Harris, 2008; Juriloff et al., 2006; Liu et al., 2005; Song et al., 2009). As a result, our understanding of molecular mechanisms involved in the formation of the lip and primary palate have lagged behind those described for formation of the secondary palate. Our identification of fully penetrant CL/P in *Esrp1*^−/−^ mice provides a new genetic model that can be exploited to further investigate mechanisms of lip development as well as to potentially define novel CL/P disease genes. It is notable that while much has been learned about signaling and transcription factors that are involved in craniofacial development, the role of alternative splicing in lip and/or palate development has been largely unexplored. Prior to our identification of CL/P in *Esrp1* KO mice, no studies have identified roles of splicing factors in CL/P. Furthermore, other than the role of a specific splice variant for *Fgfr2* (Fgfr2-IIIb), the possibility that the functions of some genes required for face and palate development are splice isoform-specific has not generally been considered. There is now firm evidence that, like transcription factors, tissue-specific splicing regulators coordinate programs of AS involving transcripts that encode proteins that function in biologically coherent pathways (Kalsotra and Cooper, 2011; Lee et al., 2018; Ule et al., 2005; Zhang et al., 2008). Thus, our studies demonstrating that *Esrp1* is required for formation of both the lip and palate indicates that gene targets of ESRP1 regulation also play essential developmental roles and that mutations in these genes may cause or predispose patients to CL/P.

*Esrp1*^−/−^ mice exhibit cleft lip associated with cleft palate (CL/P) and we first investigated whether the cleft secondary palate was an independent defect or was largely a consequence of clefting of the lip and primary palate. To discern between these possibilities and to define the epithelial cell populations that result in orofacial clefting we first crossed *Esrp1*^*flox/flox*^ mice with *Crect* transgenic mice that express Cre specifically in surface ectoderm and derivatives starting at E8.5; a stage when Esrp1 becomes restricted to ectoderm and definitive endoderm and prior to both lip and palate formation. We showed bilateral CL/P in *Esrp1*^*flox/flox*^;*Crect*^+/−^ embryos at E18.5 indicating that ablation of *Esrp1* in early surface ectoderm and derivatives underlies cleft lip and cleft palate. We considered the possibility that the cleft palate observed in *Esrp1*^−/−^ and *Esrp1*^*flox/flox*^;*Crect*^+/−^ mice embryos might be a consequence of cleft primary palate extending into the secondary palate. However, mice with compound ablation of *Esrp1* and a hypomorphic *Esrp1 Triaka* allele (*Esrp1*^*triaka/−*^ mice) had no cleft lip or primary palate but a cleft secondary palate, indicating that *Esrp1* is independently required for secondary palate (Mager et al., 2017) (Fig. 1D). These studies therefore establish that Esrp1 is required for both lip and palate development and can thus be used to further characterize mechanisms that are essential for both developmental processes. In addition, the CL/P we observed in *Esrp1*^*flox/flox*^;*Crect*^+/−^embryos confirmed that *Esrp1* ablation in early surface ectoderm is the cause of CL/P and that identification of transcriptomic changes in this cell population is key to understanding mechanisms leading to this defect.

To determine the etiology of the hypoplastic nasal and maxillary processes we examined proliferation using Ki67 staining and identified reduced staining in epithelial and mesenchymal cells of the MNP and LNP. There was also reduced Ki67 staining in the palatal shelves of *Esrp1*^−/−^ embryos at E16.5. We did not observe a notable difference in staining for apoptotic marker caspase 3, in either facial or palatine processes, indicating that a reduction in proliferation underlies the reduced size of both the nasal process as well as palatal shelves. Of note, we also found that the palatal shelves did not elevate in either *Esrp1*^−/−^ or *Esrp1*^*flox/flox*^; *Crect*^+/−^ embryos (see also Fig. 1A). However, at this stage we cannot be certain that there is also a defect in palatal elevation independent of the palatal hypoplasia that contributes to cleft secondary palate.

While we observed reduced proliferation of the MNP and LNP, to further determine whether there is also a defect in fusion of these processes contributing to cleft lip, we used a time series analysis of both WT and *Esrp1*^−/−^ embryos during several stages of lip formation. Despite the reduced proliferation, we did note several stages at which the MNP and LNP in *Esrp1*^−/−^ embryos were able to make contact and adhere, but that this did not lead to apparent fusion by E12.5, at which point fusion was complete between the LNP, MNP, and MxP in WT embryos. While we were unable to successfully complete ex vivo facial explant cultures to further verify a fusion defect between MNP, LNP, and MxP, our studies using ex vivo palatal explants demonstrated a fusion defect in *Esrp1*^−/−^ embryos as also contributing to cleft palate. Since it is believed that the mechanisms of lip and palate fusion during development are similar (Jiang et al., 2006; Ray and Niswander, 2012), we believe that the fusion defect observed in palatal explants, taken together with our histology and SEM time course also indicate a defect in fusion during lip and primary palate formation.

We carried out in depth RNA-Seq analysis to identify alternative splicing changes in *Esrp1*^−/−^ ectoderm as well as to further define differences in splicing between wild-type ectoderm and mesenchyme. Numerous changes in splicing were identified in KO vs. WT ectoderm including some events that had previously been identified in *Esrp1*^−/−^ epidermis, but the greater sequencing depth in this study identified a greater number of splicing changes than our previous studies. We first identified ESRP1 (and its paralog ESRP2) in a screen for regulators of *Fgfr2* splicing and, not surprisingly, one of the largest changes in splicing was a nearly complete switch in *Fgfr2* isoforms from Fgfr2-IIIb to Fgfr2-IIIc in *Esrp1* ablated ectoderm. This nearly complete change in splicing of Fgfr2 was in contrast to our analysis in the E18.5 epidermis that examined changes in splicing in both *Esrp1*^−/−^ and well as *Esrp1*^−/−^;*Esrp2*^−/−^ (double KO, or DKO) tissue, where there was no change in *Fgfr2* splicing unless both *Esrp1* and *Esrp2* were ablated (Bebee et al., 2015). This observation likely reflects the lower expression levels of both *Esrp1* and *Esrp2* in E12.0 surface ectoderm compared to E18.5 epidermis, including a significantly lower Esrp2 expression level compared to Esrp1 in ectoderm (see Table S3). We noted similar examples where deletion of Esrp1 alone caused greater changes in splicing in ectoderm compared to the epidermis of *Esrp1*^−/−^ mice and suspect that this is a major reason why we observe cleft palate in *Esrp1*^−/−^ mice, but not most of the other major defects previously described in *Esrp1*^−/−^;*Esrp2*^−/−^ mice. We also noted other craniofacial abnormalities in *Esrp1^−/−^;Esrp2^−/−^* mice, including mandibular defects, not seen in *Esrp1*^−/−^ mice indicating that ESRP1-regulated splicing by both paralogs also plays broader roles in craniofacial development (Bebee et al., 2015). A previous study demonstrated cleft palate, but not cleft lip, in mice in which the epithelial *Fgfr2* exon IIIb was deleted (Rice et al., 2004). However, deletion of exon IIIb in the mice did not default to splicing of exon IIIc in epithelial cells, but instead caused skipping of both exons and a frameshift that effectively resulted in no Fgfr2 expression in epithelial cells (De Moerlooze et al., 2000). In contrast, ablation of *Esrp1* induces a switch in isoforms, such that ectopic Fgfr2-IIIc in epithelial cells can still respond to Fgf ligands to sustain Fgf signaling as we demonstrated in a prior study (Rohacek et al., 2017). These observations strongly suggest that altered splicing of *Fgfr2* does not account for the cleft lip observed in *Esrp1*^−/−^ mice and is also unlikely to underlie the cleft palate.

While we identified large numbers of splicing changes in *Esrp1*^−/−^ compared to WT ectoderm, we also identified differences in splicing between WT ectoderm and WT mesenchyme. These analyses identified a large number of differences in splicing between these cell populations, which included many ESRP1 regulated events. We note that while numerous investigations have identified distinct epithelial and mesenchymal markers at the whole transcript or protein level, there remain limited examples in which large scale differences in splicing between these cell populations have been identified (Venables et al., 2013). The analysis presented here thus provides another resource to identify how different splice isoforms influence epithelial-mesenchymal crosstalk as previously described for Fgfr2 (De Moerlooze et al., 2000; Warzecha et al., 2009a).

Identification of changes in total transcript levels in *Esrp1*^−/−^ ectoderm compared to controls revealed substantial numbers of genes that were upregulated or downregulated and there was little if any overlap between these genes and those that demonstrated changes in splicing. Most striking was a coordinated decrease in the expression levels of canonical Wnts, including *Wnt9b*, as well as in *Shh*; components of two pathways that have previously been shown to be essential for lip and/or palate development (Jin et al., 2012; Lan and Jiang, 2009; Lipinski et al., 2010). These alterations are associated with corresponding downregulation of both canonical Wnt as well as Shh regulated targets in adjacent mesenchyme, suggesting a model where a reduction in a communication pathway from ectoderm to mesenchyme leads to reduced mesenchymal proliferation. Alterations in canonical Wnt signaling have been implicated in mouse models as well as human cases of syndromic and non-syndromic cleft lip and/or cleft palate (Reynolds et al., 2019), including CL/P in mice with ablation of *Wnt9b* and *Lrp6* (Carroll et al., 2005; Ferretti et al., 2011; Jin et al., 2012; Juriloff et al., 2006; Song et al., 2009). The reduced mesenchymal proliferation observed in mesenchyme of *Esrp1*^−/−^ mice is similar to that demonstrated in both *Lrp6*^−/−^ mice and *Wnt9b*^−/−^ mice, suggesting that reduced mesenchymal proliferation and failed approximation of the MNP, LNP, and MxP due to reduced Wnt signaling contribute to CL/P in our mice. However, reduced expression of Fgf ligands in ectoderm, including Fgf8 and Fgf17, was described in *Wnt9b*^−/−^ mice suggesting that reduced Fgf signaling from ectoderm to mesenchyme played a role in the etiology of CL/P in these mice. However, we did not identify decreases in the expression of these or other Fgfs in *Esrp1*^−/−^ ectoderm, but rather there was a nearly 2-fold increase in *Fgf17* transcripts. Thus, while we also propose a role for altered Wnt signaling in CL/P observed in *Esrp1*^−/−^ mice, we do not currently have evidence that downstream alterations in Fgf signaling are involved in the phenotype. In addition, *Wnt9b*^−/−^ mice did not have a defect in fusion of NP and MxP in explant cultures, whereas *Esrp1*^−/−^ mice have a fusion defect in addition to reduced mesenchymal proliferation. Hence, while the proliferation defect might be rescued by restoration of Wnt activity we suspect that alterations in other genes and pathways also contribute to the orofacial clefting defects in *Esrp1*^−/−^ mice by preventing epithelial fusion.

In the case of Shh, we noted that in addition to reductions in Gli transcription factors, there was also reduced expression of *Foxf1a*, *Foxf2*, and *Osr2* in *Esrp1*^−/−^ mesenchyme. During palatal development, a previous study showed that abrogation of Shh signaling from epithelial cell to mesenchyme through ablation of *Smo* in mesenchyme caused cleft lip associated with a reduction in these same transcription factors as well as in proliferation (Lan and Jiang, 2009). Another study showed that inhibition of Shh signaling with the inhibitor cyclopamine caused cleft lip that was also associated with a reduction of Foxf2 in mesenchyme (Everson et al., 2017). A recent study using single cell RNA-Seq to identify subpopulations of cell types present at the lambdoidal junction where the MNP, LNP, and MxP fuse characterized distinct and dynamic expression patterns in subsets of both ectodermal and mesenchymal cells during lip fusion (Li et al., 2019). This analysis revealed that while canonical Wnts were specifically expressed in the ectoderm of these processes, they were excluded from the fusion zone once these processes made contact, consistent with a lack of requirement for Wnts (at least for Wnt9b) for fusion and dissolution of the epithelial seam. This study also identified *Fgf10* among the genes that are highly expressed in mesenchymal cells at the fusion zone. During palate formation mesenchymal *Fgf10* was previously shown to induce *Shh* expression in adjacent epithelial cells via the Fgfr2-IIIb isoform and conditional ablation of *Shh* in palatal epithelial cells leads to cleft palate (Lan and Jiang, 2009). A switch in *Fgfr2* splicing in *Esrp1*^−/−^ palatal epithelium would render it unresponsive to mesenchymal *Fgf10* suggesting that the Fgfr2 splicing switch may be one factor leading to reduced Shh expression in epithelial cells of the facial processes as well as palate. In addition to Fgf10, the aforementioned single cell RNA-Seq analysis also showed a reduction in Tgfb2 in the fusion zone of both ectoderm and mesenchyme (Li et al., 2019). It is therefore tempting to speculate that the reduction in Tgfb2 observed in mesenchyme adjacent to *Esrp1* ablated ectoderm may contribute to the observed fusion defect. However, there may also be combinatorial effects of ectodermal genes that are upregulated upon *Esrp1* ablation, which includes Dkk1, Aldha3, and Sfrs2 that have been described as Wnt inhibitors.

While our results strongly implicate reductions in Wnt signaling in at least contributing to CL/P in *Esrp1*^−/−^ mice, an unresolved question remains as to how *Esrp1* ablation in ectoderm leads to coordinated changes in the expression of several canonical Wnts. Ablation of the *Pbx1* and *Pbx2* transcription factors was previously shown to lead to CL/P through reduced expression of *Wnt9b* and *Wnt3*. However, we did not identify changes in the expression of *Pbx1*, *Pbx2*, or *Pbx3* in *Esrp1* KO ectoderm to explain the associated reduction in Wnt expression. It is notable that while there is a vast literature describing numerous Wnt target genes in different contexts, there remains a limited understanding as to how the Wnt genes themselves are regulated. While there are few transcription factors among the Esrp1 regulated splicing targets in ectoderm, we did note a change in splicing of *Lef1* in ectoderm in addition to the general reduction in *Lef1* expression in mesenchyme. While this splicing event has been previously characterized, the precise changes in transcriptional activity by these Lef1 isoforms have not been well studied. We nonetheless suspect that changes in transcriptional regulation that result from *Esrp1* ablation are indirect; possibly through alterations in signaling pathways that regulate transcription factor expression and/or activity. We have previously shown that ESRP1 is concomitantly expressed as both a nuclear and cytoplasmic isoform and it is thus possible that ESRP1 might regulate RNA stability to account for some of the gene expression differences (Fagoonee et al., 2017; Yang and Carstens, 2017). However, in crosslinking-immunoprecipitation experiments we have performed in other epithelial cells we have not identified ESRP1 binding sites in canonical Wnt mRNAs (unpublished data). In any event, a major task for further studies to understand how ablation of *Esrp1* leads to CL/P will need to dissect how the loss or decrease in expression of epithelial splice isoforms in ectoderm leads to CL/P through alterations in molecular networks of splicing and transcription that are required for normal lip and palate development. The transcriptomic analysis presented here will hopefully provide a resource that can be used by the community to better understand molecular mechanisms that lead to CL/P as well as *Esrp1* regulated targets that may also merit further investigations as possible disease genes.

---

**Figure S1**. Anti-FLAG immunostaining in lip and palatal sections from Esrp1^FLAG/FLAG^ mice showing epithelial-specific expression in epithelial of the developing face and palate.

**Figure S2**. Additional examples of palatal organ cultures showing lack of dissolution of the MES in palatal shelves from *Esrp1*^−/−^ embryos compared to WT.

**Table S1**. Complete data file of alternative splicing differences identified by rMATS between wild-type ectoderm and Esrp1 KO ectoderm. Individual tabs represent splicing changes by type. Positive splicing changes were filtered for False Discovery Rate (FDR), 5% and absolute value of predicted change in Percent Spliced In (PSI) greater than or equal to 5%.

**Table S2**. Complete data file of all alternative splicing events identified by rMATS between wild-type ectoderm and wild-type mesenchyme. Tabs and filtering criteria are as indicated in legend to Table S1.

**Table S3**. Complete data file of differences in total gene expression using EdgeR between wild-type ectoderm and *Esrp1* KO ectoderm. Events were filtered for FDR < 0.05, and EdgeA>0 (corresponding to average counts per million (CPM) across all replicates to delete lowly expressed genes. Separate tabs indicate genes upregulated or downregulated in *Esrp1* KO compared to wild-type. Another tab contains unfiltered transcript per million (TPM) counts for all replicates.

**Table S4**. Complete data file of differences in total gene expression using EdgeR between wild-type mesenchyme and mesenchyme adjacent to *Esrp1* KO ectoderm. The same filtering criteria were used as for Table S3.

**Table S5.** Complete data file of differences in total gene expression using EdgeR between wild-type ectoderm and wild-type mesenchyme. The same filtering criteria were used as in the legend for Table S3 and separate tabs indicate genes that show greater expression in ectoderm or mesenchyme.

## MATERIALS and METHODS

### Mouse strains

Generation of Esrp1 KO *(Esrp1*^−/−^) and Conditional Esrp1 (*Esrp1*^*f/f*^) were described previously (Bebee et al., 2015) as was the Crect strain (Reid et al., 2011). *Axin2*^*LacZ*^ (Manuylov et al., 2008) and *TCF/Lef1:H2B-GFP* (Ferrer-Vaquer et al., 2010) strains were purchased from JAX Labs (Bar Harbor, ME). *Esrp1*^*FLAG/FLAG*^ mice were generated in mouse V6.5 ES cells by the Penn Transgenic and Chimeric mouse core facility using electroporation of an mRNA encoding Cas9, a sgRNA targeting the ATG start codon, and an oligonucleotide repair template encoding two tandem copies of the FLAG epitope tag. Relevant strains were interbred from embryo isolation and females were examined in the morning for presence of a vaginal plug, and the presence of a plug was designated E0.5. Genomic DNA for genotyping was derived from tail biopsies and genotyping was performed using standard procedures for these strains. All animal procedures and experiments were approved by the Institutional Animal Care and Use Committee (IACUC) at the University of Pennsylvania.

### Scanning Electron Microscopy

Mouse embryos were harvested at either E10.5, E11.5, or E12.5. The heads were fixed in Karnovsky's solution and a portion of the unfixed body including the tail was saved from each embryo for genotyping. Fixed head samples were dehydrated through a graded ethanol series and placed in Freon (1,1,2-Trichloro-1,2,2 Trifluoroethane) (Recycle & Reuse Industries, Mansfield, TX) for critical-point drying. The samples were mounted on aluminum stubs with clay and sputter-coated with gold in an Argon atmosphere, using a Denton Vacuum Desk II Cold Sputter Etch Unit (Denton Vacuum, Cherry Hill, NJ). The heads were then viewed under a Quanta 600 FEG Mark II scanning electron microscope (FEI, Hillsboro, OR).

### Isolation of ectoderm and mesenchyme for RNA harvest

E12.0 mouse embryos were dissected, and the ectodermal and mesenchymal tissue layers of the prominences were separated and collected for wild-type and Esrp1^−/−^ mice for a total of 4 pools of 6 to 7 mice in each category. The facial ectoderm and mesenchyme of each embryo was separated using a described surgical protocol (Li and Williams, 2013). RNA harvest of collected tissue was described previously (Bebee et al., 2015). For synthesis of cDNA, 300ng of total RNA was used for ectoderm and mesenchyme samples, oligodT primer, and SuperScript3 reverse transcriptase (Invitrogen, Carlsbad, CA).

### Real time RT-PCR and RT-PCR

Real-time RT-PCR and RT-PCR were performed as described (Bebee et al., 2015; Lee et al., 2018). Briefly, Real-time RT-PCR analysis were quantified using ImageQuant TL, version 7.0. Splicing ratios are represented as PSI for cassette exons and were normalized to RT-PCR product size. Real-time RT-PCR and RT-PCR primer sequences are listed in Supplemental Table S6 and Supplemental Table S7.

### RNA-Sequencing and data analysis

Total RNA from ectodermal and mesenchymal samples was used for RNA-Seq at the Penn Next Generation Sequencing Core (NGSC) Facility as previously described with the exception that we obtained 150 bp paired end reads (Bebee et al., 2015). The average number of read pairs was approximately 100 million read pairs per replicated. Identification of changes in alternative splicing in ectoderm samples was carried out using rMATS as previously described (Shen et al., 2014). Differential gene expression analysis was carried out by the Penn NGSC using EdgeR. Gene expression values were measured by Kallisto (v0.43.0) with mm10 gencode vM13 as the transcriptome index. The RNA-seq data will be deposited into the NCBI Gene Expression Omnibus upon manuscript publication.

### Skeletal analysis

E18.5 embryo heads were skinned, fixed in 4% Paraformaldehyde O/N. For cartilage and bone staining, E18.5 embryos were stained with Alizarin red and Alcian blue for examination of bone and cartilage structure, as previously described (Bebee et al., 2015).

### Histology and Immunofluorescence

Embryos from E10.5 to E14.5 were harvested and fixed O/N at 4% in PFA and fixed in paraffin. Paraffin sections were deparaffinized in xylene and rehydrated using a graded ethanol series. For standard histology sections were stained with H & E. For immunofluorescence, antigen retrieval was performed using unmasking solution (Vector Laboratories, Burlingame, CA) in a humidified chamber. Samples were blocked with 5% BSA and 3% sheep serum in PBST 1hour at RT. Primary antibodies were incubated overnight at 4°C. After washing with PBST, secondary antibodies were applied for 30 minutes at RT, followed by another wash and mounted with Prolong Gold antifade reagent with DAPI (Invitrogen). Images were taken using an Olympus BX43. Samples were incubated with primary antibodies against FLAG-M2 (Sigma, Cat #F1804), E-cadherin (BD Bioscience, Cat #610181), Cleaved Caspase-3 (Cell Signaling, #9664S), Ki-67 (Abcam, Ab16667), GFP (Abcam # A-11122), Beta Galactosidase (Abcam, ab9361) and secondary antibodies Rabbit IgG Alexa 488 (Life technologies, Cat #A24922), and Mouse IgG Alexa 594 (Life technologies, Cat #A24921). Quantifications of fluorescence in immunostained slides was carried out using ImageJ with comparisons between tissues corrected for area.

### Whole-mount and Section In Situ hybridization

In situ hybridization experiments were carried out as previously described (Moorman et al., 2001; Warzecha et al., 2009a) using digoxigenin-labeled riboprobes for Axin2, Lef1, Wnt9b, Shh. The signal was detected with alkaline phosphatase and then color developed with BP substrate.

### Palatal organ culture

A pair of unfused palatal shelves were dissected out from an E14 mouse embryo under a dissecting microscope and placed on autoclaved nitrocellulose membrane with the oral side facing down. The palatal shelves were correctly oriented as in vivo and gently pushed against each other to ensure contact between the medial edge epithelia. Palatal shelves on a membrane were rested on a wire grid in 12 well culture plate and cultured for 72 hours in DMEM containing 10% FBS. Tail DNA isolated from each embryo was used for genotyping. The organ culture was then processed for histology and immunofluorescent labeling.

## Acknowledgements

We thank Natoya Peart, Eric C. Liao, and Shannon Carroll for critical review of the manuscript. We thank members of the Penn Transgenic and Chimeric mouse core facility for assistance in the production of novel mouse strains. We thank Lukas F. Mager and Lester Thoo (University of Bern) for organizing neonate Triaka mice.

## Author contributions

S.L.K. performed most experiments. M.J.S, I.S., and H.D.N performed some of the experiments. H.L. and T.W provided direct training and reagents for the isolation of ectoderm and mesenchyme from E12.0 facial prominences. Z.Z. and Y.X. carried out the processing of the RNA-Seq data using rMATS for alternative splicing analysis. P. K. supervised studies of Esrp1^−/triaka^ mice that analyzed facial and palatal development. R.P. C. and S.K.L. conceived, designed the study, and wrote the manuscript.

